# Analysis of genetic networks regulating refractive eye development in collaborative cross progenitor strain mice reveals new genes and pathways underlying human myopia

**DOI:** 10.1101/530766

**Authors:** Tatiana V. Tkatchenko, Rupal L. Shah, Takayuki Nagasaki, Andrei V. Tkatchenko

**Affiliations:** Department of Ophthalmology, Columbia University, New York, New York, USA; School of Optometry & Vision Sciences, Cardiff University, Cardiff, UK; Department of Pathology and Cell Biology, Columbia University, New York, New York, USA

**Keywords:** myopia, refractive eye development, genetic networks, signaling pathways, genetic variation, RNA-seq, gene-based genome-wide association analysis, evolutionary conservation of pathways

## Abstract

**Background:** Population studies suggest that genetic factors play an important role in refractive error development; however, the precise role of genetic background and the composition of the signaling pathways underlying refractive eye development remain poorly understood.

**Methods:** Here, we analyzed normal refractive development and susceptibility to form-deprivation myopia in the eight progenitor mouse strains of the Collaborative Cross (CC). We used RNA-seq to analyze gene expression in the retinae of these mice and reconstruct genetic networks and signaling pathways underlying refractive eye development. We also utilized genome-wide gene-based association analysis to identify mouse genes and pathways associated with myopia in humans.

**Results:** Genetic background strongly influenced both baseline refractive development and susceptibility to environmentally-induced myopia. Baseline refractive errors ranged from −21.2 diopters (D) in 129S1/svlmj mice to +22.0 D in CAST/EiJ mice and represented a continuous distribution typical of a quantitative genetic trait. The extent of induced form-deprivation myopia ranged from −5.6 D in NZO/HILtJ mice to −20.0 D in CAST/EiJ mice and also followed a continuous distribution. Whole-genome (RNA-seq) gene expression profiling in retinae from CC progenitor strains identified genes whose expression level correlated with either baseline refractive error or susceptibility to myopia. Expression levels of 2,302 genes correlated with the baseline refractive state of the eye, whereas 1,917 genes correlated with susceptibility to induced myopia. Genome-wide gene-based association analysis in the CREAM and UK Biobank human cohorts revealed that 985 of the above genes were associated with myopia in humans, including 847 genes which were implicated in the development of human myopia for the first time. Although the gene sets controlling baseline refractive development and those regulating susceptibility to myopia overlapped, these two processes appeared to be controlled by largely distinct sets of genes.

**Conclusions:** Comparison with data for other animal models of myopia revealed that the genes identified in this study comprise a well-defined set of retinal signaling pathways, which are highly conserved across different vertebrate species. These results identify major signaling pathways involved in refractive eye development and provide attractive targets for the development of anti-myopia drugs.

## Background

Myopia is the most common ocular disorder worldwide [1]. The prevalence of myopia in the U.S. has increased from 25% to ~48% in the last 40 years [2-4]. The worldwide prevalence of myopia is predicted to increase from the current 25% to 50% in the next three decades [5], while the prevalence already exceeds 80% in several parts of Asia [6, 7]. Myopia often leads to serious blinding complications such as myopic maculopathy, retinal floaters, chorioretinal atrophy, retinoschisis, retinal tears, retinal detachment, and myopic macular degeneration [8-24]. It also represents a major risk factor for a number of other serious ocular pathologies such as cataract and glaucoma [9, 10, 25-27]. Because of the increasing prevalence, myopia is rapidly becoming one of the leading causes of vision loss in several parts of the world, and World Health Organization designated myopia as one of five priority health conditions [1, 8-10, 28].

Development of myopia is controlled by both environmental and genetic factors [29-32]. Although environmental factors, such as reading and nearwork, play a very important role in the development of myopia [33-36], genetic studies suggest that the impact of environmental factors on refractive development is determined by genetic variation in “myopia-susceptibility genes” [37]. The role of genetic background in refractive eye development is also supported by animal studies, which revealed that the extent of myopia experimentally induced in animal models is strongly influenced by genetic background [38-41]. Analysis of the size of ocular components in different strains of mice suggested a significant role of genetic background in the regulation of refractive eye development [42, 43]. Wong and Brown [44] also described significant differences in visual detection, pattern discrimination and visual acuity among different strains of mice. The contribution of genetic factors to myopia has been estimated to be as high as 70-80% [45-50], and human genetic mapping studies have identified over 270 chromosomal loci linked to myopia [51-54]. These loci implicate genes involved in multiple cellular and biological processes related to extracellular matrix organization, eye morphogenesis, retinal signaling, and visual perception [53, 54]. Gene expression profiling studies also showed that development of myopia is accompanied by changes in gene expression in the retina, choroid, and sclera [39, 55-62]. Moreover, these studies suggested that similar biological processes underlie refractive development in animal models and humans [63].

Although these studies revealed important roles of genetic variation and gene expression in the development of refractive errors, the relationship between genetic background, gene expression and development of refractive errors remains unexplored. Here, we systematically analyzed the role of genetic background in the regulation of retinal gene expression and signaling pathways underlying refractive eye development in eight inbred strains of mice and their association with human myopia using genome-wide gene expression profiling (RNA-seq) and gene-based genome-wide association analysis in the CREAM and UK Biobank human cohorts. We found that both baseline refractive state of the eye and susceptibility to myopia are inherited as quantitative traits, demonstrating strong dependence on genetic background. Furthermore, genetic background strongly influenced expression of genes in the retina and modulated a well-defined set of signaling pathways highly conserved in chickens, mice, monkeys, and humans. Our data suggest that refractive eye development is regulated by hundreds to thousands of genes across ocular tissues and point to high evolutionary conservation of signaling pathways underlying refractive development across vertebrate species.

## Methods

### Ethics statement

Mice were obtained from the Jackson Laboratory (Bar Harbor, ME) and were maintained as an in-house breeding colony. All procedures adhered to the Association for Research in Vision and Ophthalmology (ARVO) statement on the use of animals in ophthalmic and vision research and were approved by the Columbia University Institutional Animal Care and Use Committee. Animals were anesthetized via intraperitoneal injection of ketamine (90 mg/kg) and xylazine (10 mg/kg) and were euthanized using CO_2_ followed by cervical dislocation.

All human studies were approved by the relevant institutional review boards and/or medical ethics committees and conducted according to the Declaration of Helsinki. All CREAM participants provided written informed consent. The UK Biobank received ethical approval from the National Health Service National Research Ethics Service (reference 11/NW/0382).

### Analysis of refractive state of the eyes in mice

To examine the effect of genetic background on the refractive state of the eye in mice, we analyzed baseline refractive errors in the eight strains of mice comprising Collaborative Cross. The refractive state of both left and right eyes was determined on alert animals at P40 using an automated eccentric infrared photorefractor as previously described [64, 65]. The animal to be refracted was immobilized using a restraining platform, and each eye was refracted along the optical axis in dim room light (< 1 lux), 20-30 min. after instilling 1% tropicamide ophthalmic solution (Alcon Laboratories) to ensure mydriasis and cycloplegia. Five independent measurement series (~300-600 measurements each) were taken for each eye. The measurements were automatically acquired by the photorefractor every 16 msec. Each successful measurement series (i.e., Purkinje image in the center of the pupil and stable refractive error for at least 5 sec.) was marked by a green LED flash, which was registered by the photorefractor software. Sixty individual measurements from each series, immediately preceding the green LED flash, were combined, and a total of 300 measurements (60 measurements x 5 series = 300 measurements) were collected for each eye. Data for the left and right eyes were combine (600 measurements total) to calculate mean refractive error and standard deviation for each animal.

### Analysis of form-deprivation myopia in mice

To examine the effect of genetic background on susceptibility to environmentally induced myopia in mice, we analyzed the extent of myopia induced by the diffuser-imposed retinal image degradation (visual form deprivation) in the eight strains of mice comprising Collaborative Cross. Visual input was degraded in one of the eyes by applying plastic diffusers, and refractive development of the treated eye was compared to that of the contralateral eye, which was not treated with a diffuser, as previously described [66, 67]. Diffusers represented low-pass optical filters, which degraded the image projected onto the retina by removing high spatial frequency details. Frosted hemispherical plastic diffusers were hand-made from zero power rigid contact lenses made from OP3 plastic (diameter = 7.0 mm, base curve = 7.0 mm; Lens.com). Lenses were frosted using a fine sandpaper and inserted into a 3D-printed plastic frames (Proto Labs). On the first day of the experiment (P24), animals were anesthetized via intraperitoneal injection of ketamine and xylazine, and frames with diffusers were attached to the skin surrounding the right eye with six stitches using size 5-0 ETHILON™ microsurgical sutures (Ethicon) and reinforced with Vetbond™ glue (3M Animal Care Products) (the left eye served as a control). Toenails were covered with adhesive tape to prevent mice from removing the diffusers. Animals recovered on a warming pad and were then housed under low-intensity constant light in transparent plastic cages for the duration of the experiment as previously described [66, 67]. Following 21 days of visual form deprivation (from P24 through P45), diffusers were removed and refractive status of both treated and control eyes was assessed using an automated eccentric infrared photorefractor as previously described [64, 65]. The interocular difference in refraction between the treated and contralateral control eye served as an indication of the extent of induced myopia.

### RNA extraction and RNA-seq

Animals were euthanized following an IACUC-approved protocol. Eyes were enucleated, the retinae were dissected from the enucleated eyes and the choroid/RPE removed. The retinae were washed in RNAlater (Thermo Fisher Scientific) for 5 min., frozen in liquid nitrogen, and stored at −80°C until processed for this study. To isolate RNA, tissue samples were homogenized at 4°C in a lysis buffer using Bead Ruptor 24 tissue homogenizer (Omni). Total RNA was extracted from each tissue sample using miRNAeasy mini kit (QIAGEN) following the manufacturer’s protocol. The integrity of RNA was confirmed by analyzing 260/280 nm ratios (Ratio_260/280_ = 2.11-2.13) on a Nanodrop (Thermo Scientific) and the RNA Integrity Number (RIN = 9.0-10.0) using Agilent Bioanalyzer. Illumina sequencing libraries were constructed from 1 μg of total RNA using the TruSeq Stranded Total RNA LT kit with the Ribo-Zero Gold ribosomal RNA depletion module (Illumina). Each library contained a specific index (barcode) and were pooled at equal concentrations using the randomized complete block (RCB) experimental design before sequencing on Illumina HiSeq 2500 sequencing system. The number of libraries per multiplexed sample was adjusted to ensure sequencing depth of ~70 million reads per library (paired-end, 2 × 100 nucleotides). The actual sequencing depth was 76,773,554 ± 7,832,271 with read quality score 34.5 ± 0.4.

### Post-sequencing RNA-seq data validation and analysis

The FASTQ raw data files generated by the Illumina sequencing system were imported into Partek Flow software package (version 7.0.18.1210, Partek), libraries were separated based on their barcodes, adapters were trimmed and remaining sequences were subjected to pre-alignment quality control using Partek Flow pre-alignment QA/QC module. After the assessment of various quality metrics, bases with the quality score < 34 were removed (≤ 5 bases) from each end. Sequencing reads were then mapped to the mouse reference genome Genome Reference Consortium Mouse Build 38 (GRCm38/mm10, NCBI) using the STAR aligner (version 2.5.2b) resulting in 95.0 ± 0.4% mapped reads per library, which covered 35.4 ± 1.0% of the genome. Aligned reads were quantified to transcriptome using Partek E/M annotation model and the NCBI’s RefSeq Transcripts 80 annotation file to determine read counts per gene/genomic region. The generated read counts were normalized by the total read count and subjected to the analysis of variance (ANOVA) to detect genes whose expression correlates with either refractive error or susceptibility to myopia. Differentially expressed transcripts were identified using a P-value threshold of 0.05 adjusted for genome-wide statistical significance using Storey’s q-value algorithm [68]. To identify sets of genes with coordinate expression, differentially expressed transcripts were clustered using Partek Flow hierarchical clustering module using average linkage for the cluster distance metric and Euclidean distance metric to determine the distance between data points. Each RNA-seq sample was analyzed as a biological replicate, thus, resulting in three biological replicates per strain.

### Gene ontology analysis and identification of canonical signaling pathways

To identify biological functions (gene ontology categories), which were significantly affected by the genes whose expression correlated with either baseline refractive errors or susceptibility to myopia, we used the database for annotation, visualization and integrated discovery (DAVID) version 6.8 [69] and GOplot R package [70]. DAVID uses a powerful gene-enrichment algorithm and DAVID Gene Concept database to identify biological functions (gene ontology categories) affected by differential genes, while GOplot integrates gene ontology information with gene expression information and predicts the effect of gene expression changes on biological processes. DAVID uses a modified Fisher’s exact test (EASE score) with a P-value threshold of 0.05 to estimate statistical significance of enrichment for specific gene ontology categories. IPA Pathways Activity Analysis module (QIAGEN) was used to identify canonical pathways effected by the genes involved in baseline refractive eye development or regulating susceptibility to myopia, and predict the effect of gene expression changes in different strains on specific pathways. The activation z-score was employed in the IPA Pathways Activity Analysis module to predict activation or suppression of the canonical pathways. The z-score algorithm is designed to reduce the chance that random data will generate significant predictions. The z-score provides an estimate of statistical quantity of change for each pathway found to be statistically significantly affected by the changes in gene expression. The significance values for the canonical pathways were calculated by the right-tailed Fisher’s exact test. The significance indicates the probability of association of molecules from a dataset with the canonical pathway by random chance alone. Pathways Activity Analysis module determines if canonical pathways, including functional end-points, are activated or suppressed based on the gene expression data in a dataset. Once statistically significant canonical pathways were identified, we subjected the datasets to the Core Functional Analysis in IPA to compare the pathways and identify key similarities and differences in the canonical pathways underlying baseline refractive development and susceptibility to myopia.

### Identification of candidate genes for human myopia within known myopia QTLs

To identify candidate genes for myopia in the QTLs previously found to be linked to human myopia, we compared the genes that we found to be involved in refractive eye development in mice with a list of genes located within human myopia QTLs. We first compiled a list of all SNPs or markers exhibiting statistically significant association with myopia in the human linkage or GWAS studies. LDlink’s LDmatrix tool (National Cancer Institute) was used to identify SNPs in linkage disequilibrium and identify overlapping chromosomal loci. We then used UCSC Table Browser to extract all genes located within critical chromosomal regions identified by the human linkage studies or within 200 kb (±200 kb) of the SNPs found by GWAS. The list of genes located within human QTLs was compared with the list of genes that we found to be associated with either baseline refractive errors or susceptibility to myopia in mice using Partek Genomics Suite (Partek). The statistical significance of the overlaps was estimated using probabilities associated with the hypergeometric distribution using Bioconductor software package GeneOverlap version 1.14.0 and associated functions.

### Identification of genes associated with refractive error in the human population using gene-based genome-wide association analysis

To identify genes associated with the development of refractive errors in humans among the genes whose expression correlated with refractive eye development in mice, human homologs of candidate mouse genes were examined for association with refractive error in the international genome-wide association study (GWAS) of refractive error carried out by the Consortium for Refractive Error and Myopia (CREAM) [54] and the UK Biobank Eye and Vision consortium sample [71] using the Multi-marker Analysis of GenoMic Annotation (MAGMA) [72]. Human homologs of candidate mouse genes were obtained from Ensembl BioMart and were mapped according to gene definitions in the NCBI Entrez Gene database. Genes were defined according to NCBI build 37 (hg19/GRCh37) coordinates with 200kb flanking regions appended to the transcription start/stop sites. LD patterns were estimated by MAGMA using the 1000 Genomes Phase 1, version 3 European ancestry reference panel. As summary statistics were used as input, MAGMA gene-based analysis was performed using the default “snp-wise=mean” model.

The CREAM sample included 148,485 individuals of European ancestry from 28 cohorts and 11,935 individuals of Asian ancestry from eight studies. All participants included in this analysis from CREAM were 25 years of age or older.

UK Biobank is a large prospective study following the health and wellbeing of approximately 500,000 UK residents aged between 40 and 69 years-old at the baseline recruitment visit (during the period 2006–2010). 130,521 participants had non-cycloplegic autorefraction performed for at least one eye using the Tomey RC 5000 autorefractor-keratometer (Tomey Corp., Nagoya, Japan), with up to ten measurements taken for each eye. After the exclusion of unreliable readings, 130,459 participants had measures for refractive astigmatism and spherical equivalent refractive error.

Participants with conditions that might alter refraction, such as cataract surgery, laser refractive procedures, retinal detachment surgery, keratoconus, or ocular or systemic syndromes were excluded from the analyses. Refractive error was represented by measurements of refraction and analyzed as spherical equivalent (SphE = spherical refractive error +1/2-cylinder refractive error).

## Results

### Genetic background modulates baseline refractive eye development and susceptibility to myopia in mice

Genetic background was shown to influence the size of ocular components in mice. To investigate whether genetic differences between mice would have an impact on the baseline refractive state of the eye and susceptibility to myopia, we analyzed baseline refractive development and susceptibility to form-deprivation myopia in eight inbred strains of mice, which served as founder strains for the Collaborative Cross (CC) [73], i.e., 129S1/svlmj, A/J, C57BL/6J, CAST/EiJ, NOD/ShiLtJ, NZO/HlLtJ, PWK/PhJ, and WSB/EiJ mice.

We first measured baseline refractive errors in all eight strains at P40, i.e., when baseline refractions reach a plateau in mice [64]. We found that C57BL/6J mice were emmetropic on average (+0.3 ± 0.9 D). CAST/EiJ, NZO/HlLtJ, PWK/PhJ, and WSB/EiJ mice exhibited various degrees of hyperopia ranging from +10.6±2.2 D to +22±4.0 D, whereas A/J, NOD/ShiLtJ, and 129S1/svlmj mice developed various degrees of myopia ranging from −3.5±3.6 D to −21.2±3.9 D (Additional file 1: Table S1, Fig. 1A). The differences between the strains were statistically significant as revealed by ANOVA (F(7, 145) = 429.76, *P* < 0.00001). More importantly, the distribution of refractive errors in the CC mice was continuous, suggesting that refractive state of the eye in mice is inherited as a quantitative trait.

**Fig 1.**
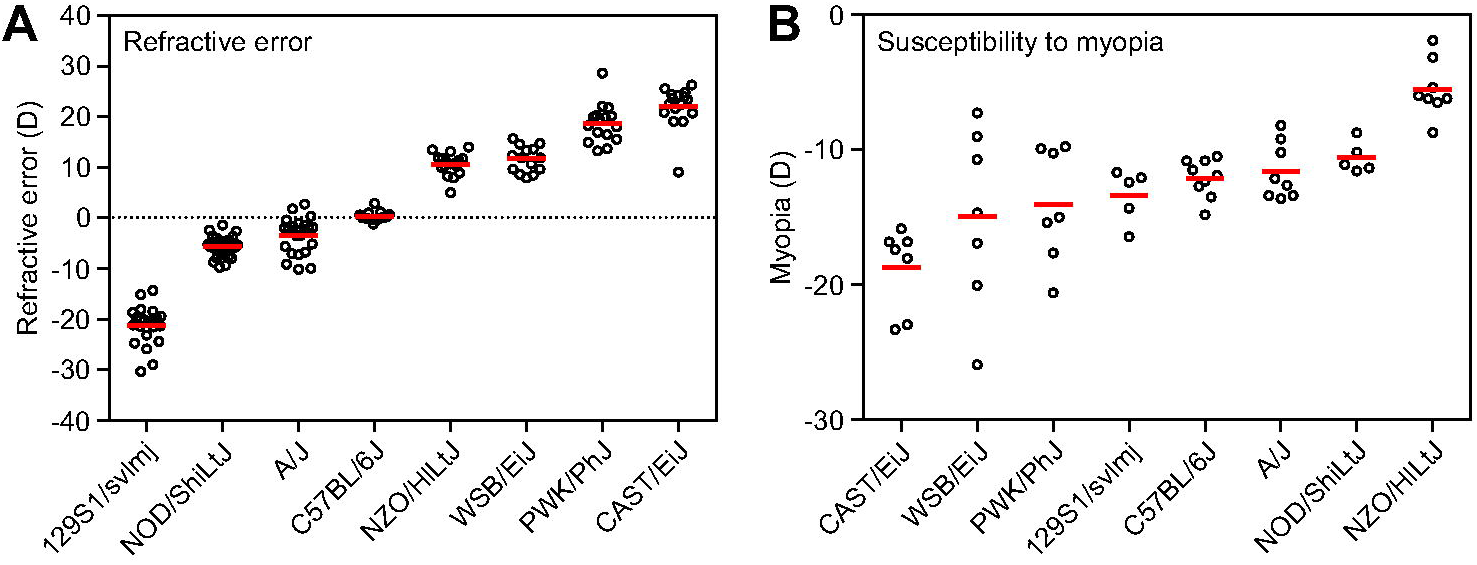
Genetic background modulates refractive eye development and susceptibility to myopia in mice. **a** Refractive error is inherited as a quantitative trait in the founder strains of Collaborative Cross. Baseline refractive errors at P40 range from highly myopic to highly hyperopic depending on the genetic background in different strains. Horizontal red lines show mean refractive errors for each strain, while each dot corresponds to mean refractive errors of individual animals. **b** Susceptibility to form-deprivation myopia is inherited as a quantitative trait in the founder strains of Collaborative Cross. The extent of myopia induced by 21 days of visual form deprivation in different strains ranged from −5.5 ± 2.1 D in NZO/HlLtJ mice to −18.7 ± 3.1 D in CAST/EiJ mice. Horizontal red lines identify means of induced myopia for each strain, while each dot represents a mean interocular difference between deprived eye and control contralateral eye for individual animals.

We then analyzed susceptibility to myopia in the same strains of mice by evaluating the extent of induced form-deprivation myopia (Additional file 2: Table S2, Fig. 1B). Susceptibility to form-deprivation myopia was estimated by applying a diffuser to one eye at P24 and comparing the extent of induced myopia in the form-deprived eye versus the contralateral control eye after 21 days of treatment. We found large differences in susceptibility to induced myopia between the strains (Additional file 2: Table S2, Fig 1B), which ranged from −5.5 ± 2.1 D in NZO/HlLtJ mice to −18.7 ± 3.1 D in CAST/EiJ mice. Other strains occupied intermediate positions between NZO/HlLtJ and CAST/EiJ mice and differences between the strains in the extent of induced myopia were statistically significant as revealed by ANOVA (F(7, 48) = 9.8, *P* < 0.00001). The distribution of induced refractive errors was continuous, similar to baseline refractive errors, suggesting that susceptibility to myopia was also inherited as a quantitative trait in mice. Spearman’s rank-order correlation analysis showed that there was no statistically significant correlation between baseline refractive errors and the extent of induced myopia (*r_s_* = −0.60, *P* = 0.12). Collectively, these data suggest that genetic background plays important role in baseline refractive development and susceptibility to environmentally induced myopia in mice and that both baseline refractive error and susceptibility to myopia are inherited as quantitative traits.

### Large number of genes are involved in regulation of baseline refractive error in mice via multiple retinal biological processes and signaling pathways

To identify retinal genes influencing baseline refractive eye development in mice, we used RNA-seq to analyze gene expression in the retina of eight CC strains at P28 (an age when refractive development is progressing towards its stable plateau) (Fig 2). We found that expression of 2,302 retinal genes strongly correlated with the baseline refractive state (Additional file 3: Table S3, Fig 2A). Genes were organized in two distinct clusters. Expression of 793 genes comprising the first cluster was positively correlated with hyperopia, i.e., expression was increased in the strains with positive refractive errors and decreased in the strains with negative refractive errors. Conversely, expression of 1,509 genes comprising the second cluster was positively correlated with myopia, i.e., expression of these genes was increased in the mouse strains with negative refractive errors and decreased in the strains with positive refractive errors. We observed a clear transition from the “hyperopic” gene expression pattern in CAST/EiJ and PWK/PhJ mice with highly hyperopic refractive errors to the “myopic” gene expression pattern in 129S1/svlmj mice with highly myopic refractive errors. Other strains occupied intermediate positions between these two extremes and exhibited transitional patterns of gene expression, which correlated with average baseline refractive errors in these strains.

**Fig 2.**
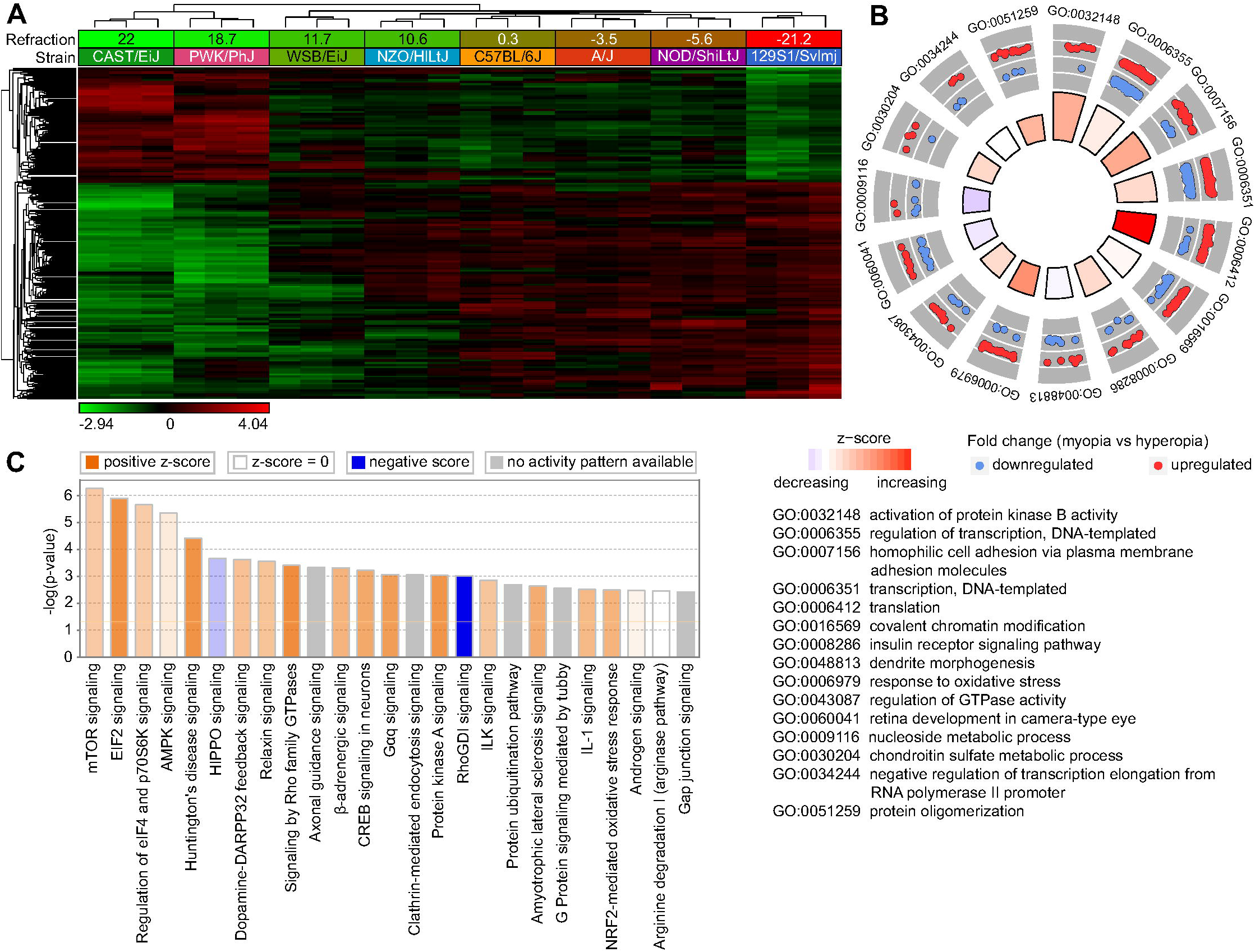
Baseline refractive eye development in mice is regulated by a large number of genes via multiple retinal signaling pathways. **a** Expression of 2,302 retinal genes correlates with baseline refractive error in mice. Hierarchical clustering results show that genes, whose expression correlates with refractive error in the founder strains of Collaborative Cross, are organized in two clusters, i.e., one (top) cluster exhibiting increased expression in the highly hyperopic mice, and the second (bottom) cluster showing increased expression in the myopic mice. **b** Top fifteen biological processes affected by the genes whose expression correlates with the baseline refractive errors in the founder strains of Collaborative Cross. Outer circle shows gene ontology IDs for the biological processes; second circle shows up- or down-regulated genes in the myopic mice versus hyperopic mice; inner circle shows activation or suppression of the corresponding biological processes, while the size of the sector corresponds to statistical significance (larger sectors correspond to smaller *P*-values). **c** Top twenty-five canonical pathways affected by the genes whose expression correlates with the baseline refractive errors in the founder strains of Collaborative Cross. Horizontal yellow line indicates *P* = 0.05. Z-score shows activation or suppression of the corresponding pathways.

Gene ontology analysis revealed that the 2,302 genes whose expression correlated with baseline refractive error were associated with 116 biological processes, 77 cellular components, and 69 molecular functions in the retina (Additional file 4: Table S4). Biological processes involved in the regulation of baseline refractive development ranged from regulation of neurogenesis and neuron migration to regulation of DNA methylation, visual perception, and synaptic vesicle endocytosis. Figure 2B shows the top 15 biological processes associated with these genes, including regulation of protein kinase B, regulation of transcription and translation, covalent chromatin modification, insulin receptor signaling, dendrite morphogenesis, and response to oxidative stress, among others. Genes underlying baseline refractive development were also associated with multiple canonical signaling pathways in the retina (Additional file 5: Table S5, Fig 2C). Negative refractive errors were associated with activation of mTOR, EIF2, AMPK, β-adrenergic, and dopamine-DARPP32 feedback signaling pathways and suppression of HIPPO and RhoGDI signaling pathways, among others. Taken together, these data suggest that refractive eye development is regulated by a large number of genes and pathways. In summary, the development of hyperopic and myopic refractive errors was associated with specific patterns of gene expression, and the activation or suppression of many retinal signaling pathways.

### Large number of genes are involved in regulation of susceptibility to myopia in mice via multiple retinal biological processes and signaling pathways

We found that genetic background influences susceptibility to experimentally induced myopia in mice and that susceptibility to myopia appears to be controlled as a quantitative trait by multiple genes (Fig 1B). To identify genes underlying inter-strain differences in susceptibility to myopia in mice, we analyzed gene expression in the retina of the eight CC strains upon induction of form-deprivation myopia at the whole-genome level, using RNA-seq (Fig 3). We found that expression of 1,917 genes strongly correlated with the extent of form-deprivation myopia in different strains of mice (Additional file 6: Table S6, Fig 3A). Similar to what we found with baseline refractive errors, these genes were organized in two clusters. Expression of 643 genes comprising the first cluster was positively correlated with the susceptibility to myopia, whereas expression of 1,274 genes in the second cluster was negatively correlated with the susceptibility to myopia. We found that there was a transition from a “high susceptibility” gene expression pattern in CAST/EiJ mice, which developed −18.7 ± 3.1 D of myopia, to a “low susceptibility” gene expression pattern in NZO/HlLtJ mice, in which only −5.5 ± 2.1 D of myopia was induced by visual form deprivation. Other strains exhibited transitional gene expression patterns, which correlated with the extent of induced myopia.

**Fig 3.**
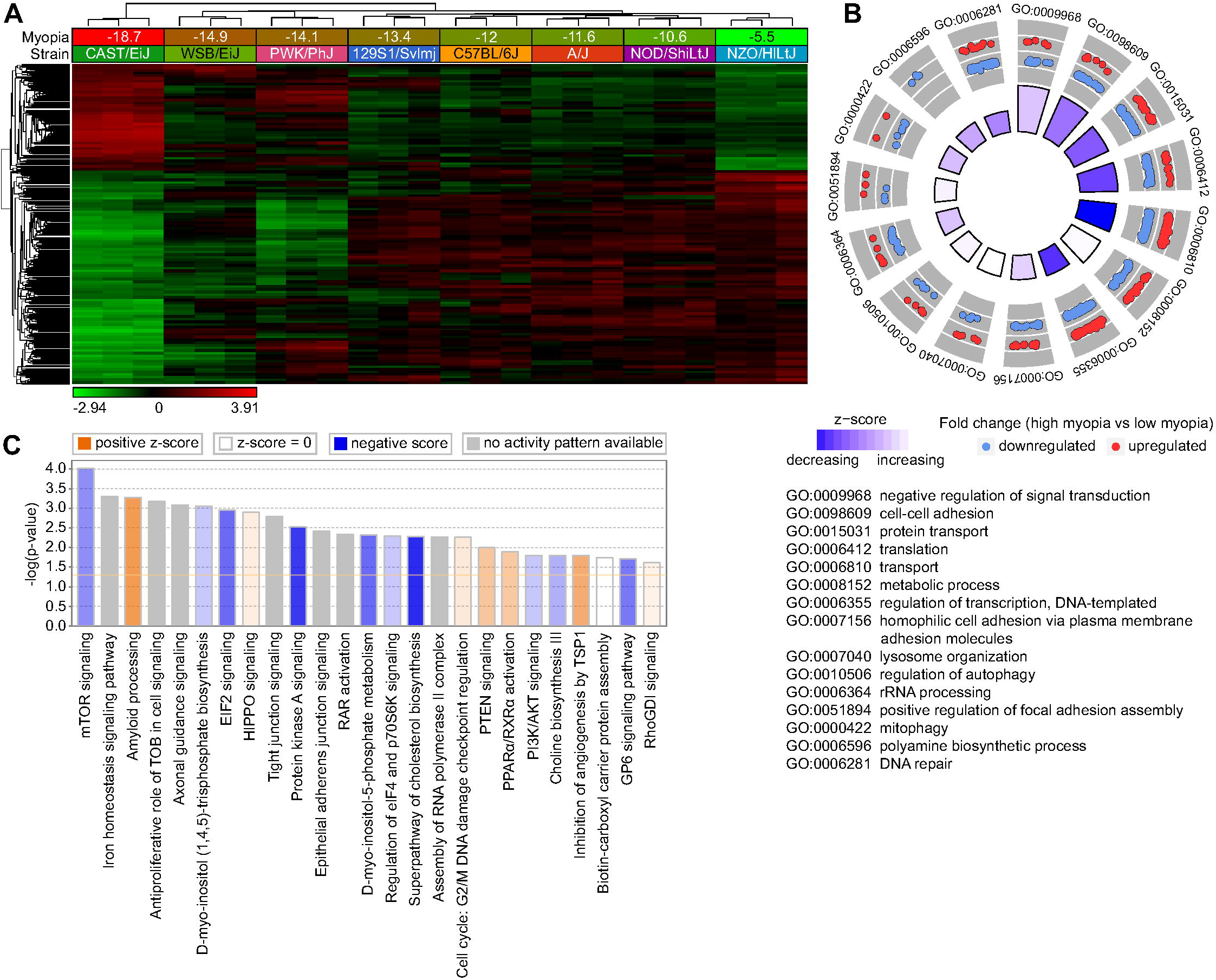
Susceptibility to myopia in mice is regulated by a large number of genes via multiple retinal signaling pathways. **a** Expression of 1,917 retinal genes correlates with susceptibility to form deprivation myopia in mice. Hierarchical clustering results show that genes, whose expression correlates with susceptibility to myopia in the founder strains of Collaborative Cross, are organized in two clusters, i.e., one (top) cluster exhibiting increased expression in mice with high susceptibility to myopia, and the second (bottom) cluster showing increased expression in mice with low susceptibility to myopia. **b** Top fifteen biological processes affected by the genes whose expression correlates with susceptibility to myopia in the founder strains of Collaborative Cross. Outer circle shows gene ontology IDs for the biological processes; second circle shows up- or down-regulated genes in mice with high susceptibility to myopia versus mice with low susceptibility to myopia; inner circle shows activation or suppression of the corresponding biological processes, while the size of the sector corresponds to statistical significance (larger sectors correspond to smaller *P*-values). **c** Top twenty-five canonical pathways affected by the genes whose expression correlates with susceptibility to myopia in the founder strains of Collaborative Cross. Horizontal yellow line indicates *P* = 0.05. Z-score shows activation or suppression of the corresponding pathways.

Gene ontology analysis suggested that 55 biological processes, 61 cellular components, and 41 molecular functions were associated with the 1,917 genes correlated with susceptibility to myopia (Additional file 7: Table S7). Figure 3B shows the top 15 biological processes involved in the regulation of susceptibility to myopia, including regulation of signal transduction, cell-cell adhesion, transcription, translation, protein transport, and lysosome organization, among others. Analysis of the canonical pathways influenced by the genes correlated with susceptibility to myopia revealed that increased susceptibility to myopia was associated with suppression of mTOR signaling, EIF2 signaling, protein kinase A signaling, D-myo-inositol-5-phosphate metabolism, cholesterol and choline biosynthesis, as well as with activation of amyloid processing, HIPPO signaling, PTEN signaling, and PPARα/RXRα signaling pathways (Additional file 8: Table S8, Fig 3C). Collectively, these data implicate an elaborate retinal genetic network and multiple signaling pathways in the regulation of susceptibility to myopia in mice.

### Baseline refractive eye development and susceptibility to myopia in mice are regulated via overlapping but largely distinct retinal genetic networks

To estimate the relative contribution of genes whose expression level correlated with baseline refractive error versus susceptibility to myopia, we analyzed the overlap between the two gene sets (Fig 4). We found that 714 genes were correlated with both baseline refractive development and susceptibility to myopia (Additional file 9: Table S9, Fig 4A). Gene ontology analysis revealed that these 714 genes were associated with 24 biological processes, 24 cellular components, and 14 molecular functions (Additional file 10: Table S10). Figure 4B shows the top 15 biological processes for the overlapping genes. Interestingly, the majority of these biological processes were suppressed in animals with high susceptibility to myopia (lower panel) and were activated in animals with negative baseline refractive errors (upper panel). A similar trend was observed when we compared canonical signaling pathways affected by the overlapping genes (Additional file 11: Table S11, Fig 4C). Signaling pathways, which were activated in mice with highly negative baseline refractive errors, were suppressed in mice with high susceptibility to myopia, and vice versa. The top pathways associated with both baseline refractive development and susceptibility to myopia were EIF2 signaling, protein kinase A signaling, regulation of eIF4 and p70S6K signaling, mTOR pathway, HIPPO pathway, and axonal guidance signaling. Figure 5 shows a summary of all signaling pathways correlated with baseline refractive eye development and susceptibility to myopia. In addition to the pathways listed above, a number of other pathways were also implicated in both baseline refractive development and susceptibility to myopia, including GP6 signaling pathway, melatonin signaling, RhoGDI signaling, PTEN signaling, opioid signaling pathway, PPARα/RXRα activation, PI3K/AKT signaling, estrogen receptor signaling, and tight junction signaling. Conversely, synaptic long-term potentiation, α-adrenergic and β-adrenergic signaling, androgen and aldosterone signaling, ephrin receptor signaling, relaxin signaling, dopamine-DARPP32 feedback signaling, dopamine receptor signaling, eNOS and nNOS signaling, somatostatin receptor 2 signaling, neurotrophin/TRK signaling, protein ubiquitination pathway, gap junction signaling, phototransduction pathway, and several other pathways were associated with baseline refractive development but not susceptibility to myopia. The amyloid processing, IGF-1 signaling, DNA methylation and transcriptional repression signaling, epithelial adherens junction signaling, iron homeostasis signaling pathway, RAR activation, RAN signaling, and several other pathways were associated with susceptibility to myopia but not baseline refractive error. Thus, these data suggest that baseline refractive development and susceptibility to myopia are regulated by genetic networks with considerable overlap; however, the two genetic networks have substantial unique components, which may independently regulate either baseline refractive development or susceptibility to environmentally induced myopia.

**Fig 4.**
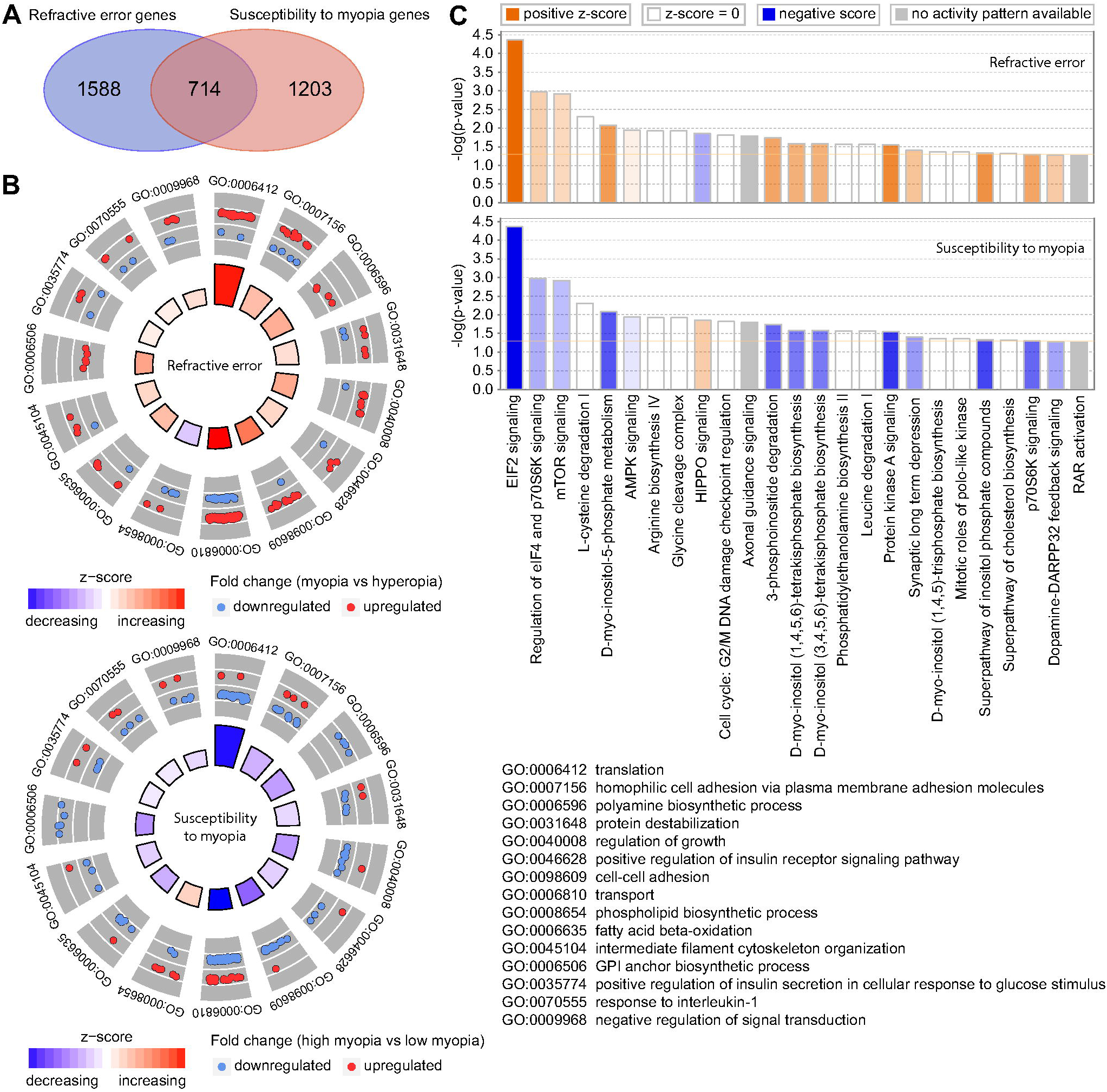
Baseline refractive eye development and susceptibility to myopia in mice are regulated via overlapping pathways. **a** Venn diagram showing substantial overlap between genes underlying baseline refractive eye development and genes regulating susceptibility to myopia in Collaborative Cross progenitor strain mice. **b** Top fifteen biological processes affected by the 714 genes associated with both baseline refractive eye development (top panel) and susceptibility to myopia in mice (bottom panel). Outer circle shows gene ontology IDs for the biological processes; second circle shows up- or down-regulated genes in the myopic mice versus hyperopic mice (top panel), or in mice with high susceptibility to form-deprivation myopia versus mice with low susceptibility to myopia (bottom panel); inner circle shows activation or suppression of the corresponding biological processes, while the size of the sector corresponds to statistical significance (larger sectors correspond to smaller *P*-values). **c** Top twenty-five canonical pathways affected by the 714 genes involved in the regulation of both baseline refractive eye development (top panel) and susceptibility to myopia (bottom panel) in the founder strains of Collaborative Cross. Horizontal yellow line indicates *P* = 0.05. Z-score shows activation or suppression of the corresponding pathways.

**Fig 5.**
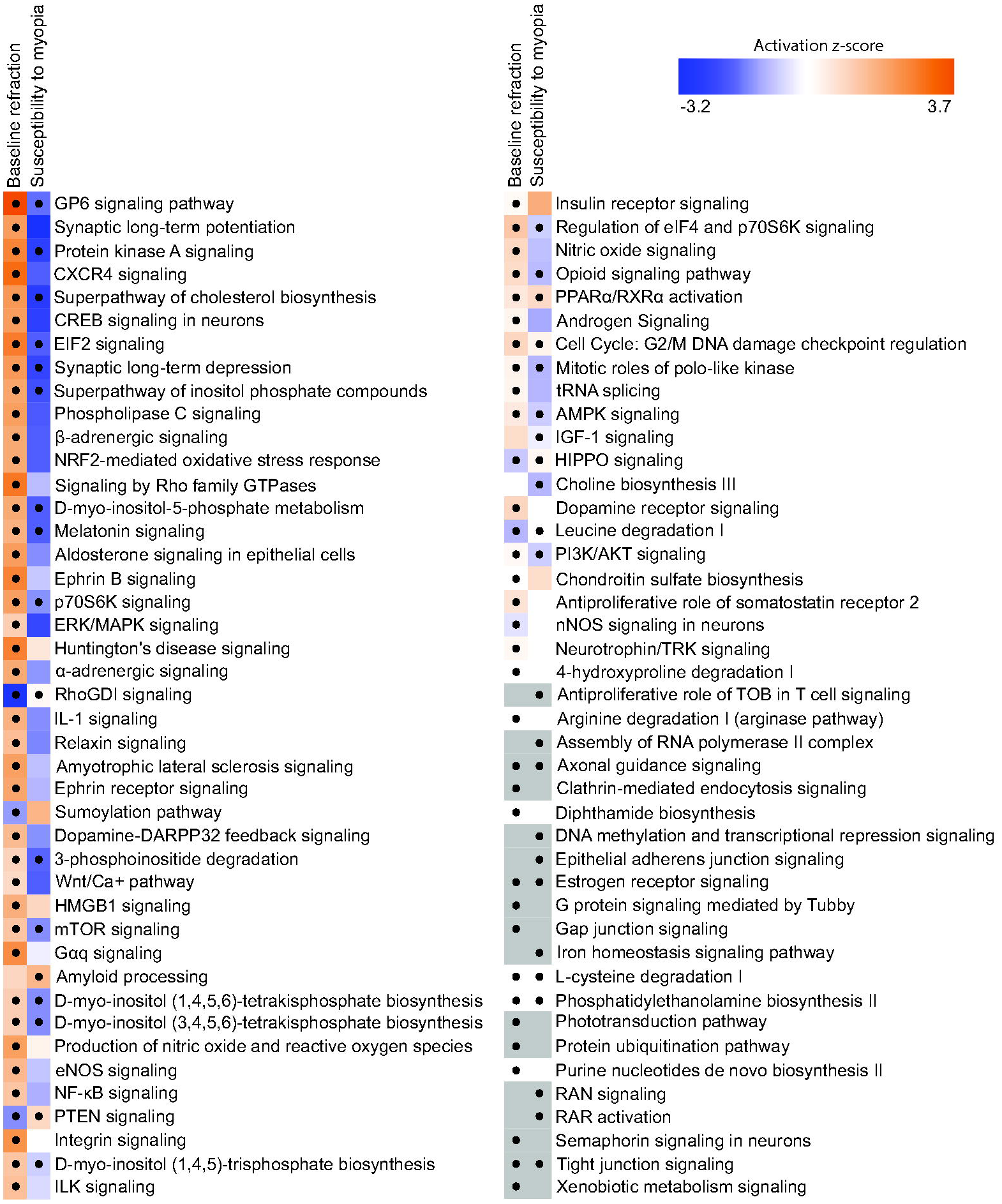
Summary of signaling pathways involved in regulation of baseline refractive eye development and susceptibility to myopia. Heatmap showing all statistically significant canonical pathways affected by the genes involved in baseline refractive development and the genes influencing susceptibility to myopia. Z-score shows activation or suppression of the corresponding pathways. The dots show statistical significance (*P* < 0.05).

### Many genes regulating baseline refractive eye development or susceptibility to myopia in mice are localized within chromosomal loci linked to human myopia

Genes comprising genetic networks underlying important developmental and physiological processes often harbor mutations causing human diseases. Therefore, to identify genes associated with susceptibility to myopia in humans, we analyzed the overlap between the genes that we found to be associated with either baseline refractive development or susceptibility to myopia in mice and genes located within human quantitative trait loci (QTLs) linked to myopia (Additional file 12: Table S12, Fig 6, 7 and 8). We found that 90 genes whose expression correlated with baseline refractive errors, 51 genes whose expression correlated with susceptibility to myopia, and 23 genes whose expression correlated with both baseline refractive errors and susceptibility to myopia in mice were localized within human QTLs linked to myopia (Additional file 13: Table S13, Additional file 14: Table S14, Additional file 15: Table S15, Fig 6). GeneOverlap analysis (Fig 6) revealed that the overlaps for the genes involved in baseline refractive development and genes involved in the regulation of susceptibility to myopia were highly significant (OR = 2.75, *P* = 3.3 × 10^−18^; OR = 2.03, *P* = 1.2 × 10^−07^; respectively). The overlap between the genes located within human QTLs and genes associated with both baseline refractive development and susceptibility to myopia in mice was weaker (OR = 1.62, *P* = 0.022); however, overall, these data suggest a functional association between genes associated with refractive eye development in mice and genes causing myopia in humans. GeneOverlap analysis revealed that a total of 164 mouse genes were located within 109 human QTLs, producing 1.5 candidate genes per QTL (Additional file 16: Table S16, Fig 7 and 8).

**Fig 6.**
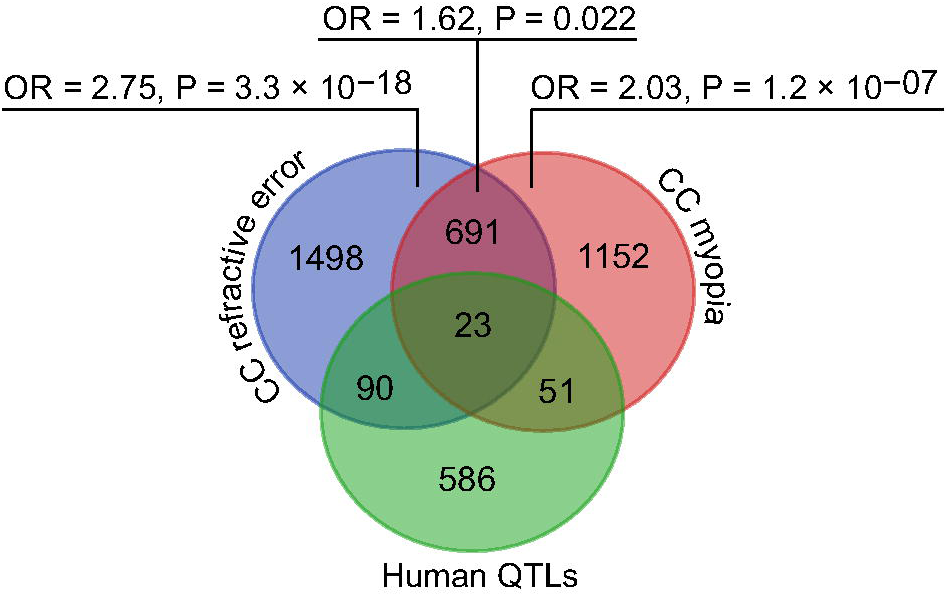
Genes localized within human myopia QTLs show functional overlap with genes underlying baseline refractive eye development and susceptibility to myopia in mice. 750 candidate genes localized within 279 human myopia QTLs exhibit statistically significant overlap with both genes involved in baseline refractive eye development and genes regulating susceptibility to myopia in Collaborative Cross progenitor strain mice.

**Fig 7.**
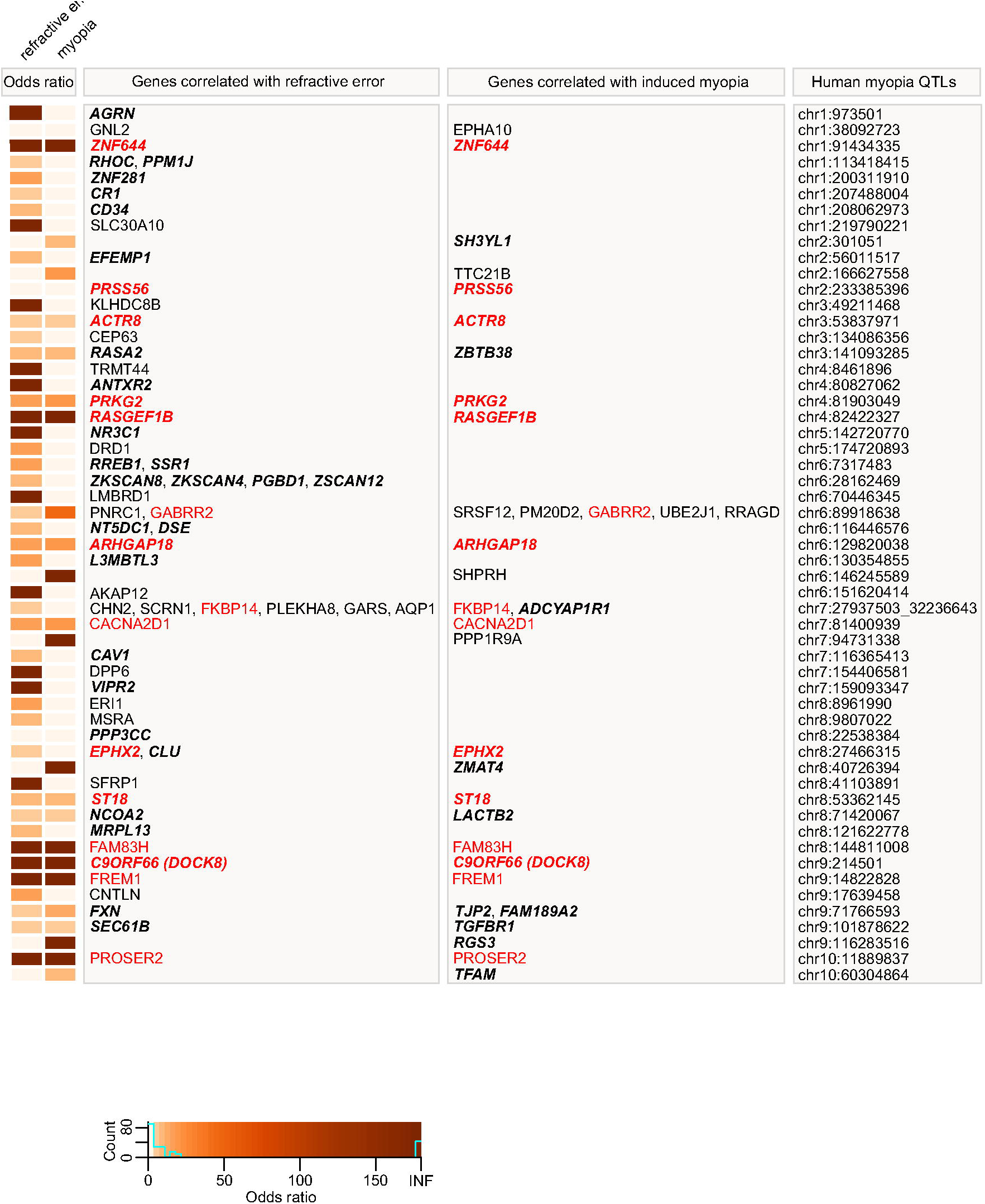
Genes underlying baseline refractive eye development and susceptibility to myopia in mice are localized in human QTLs associated with myopia (part 1). Heatmap depicting genes and odds ratios for the overlaps between 109 human myopia QTLs and genes whose expression correlates with either baseline refractive errors, susceptibility to form deprivation myopia, or both in mice. Colors indicate odds ratios. Bold italic identifies genes found to be associated with refractive error in UK Biobank, CREAM, or both human samples by the gene-based genome-wide association analysis. Red identifies genes exhibiting correlation with both baseline refractive development and susceptibility to myopia in mice.

**Fig 8.**
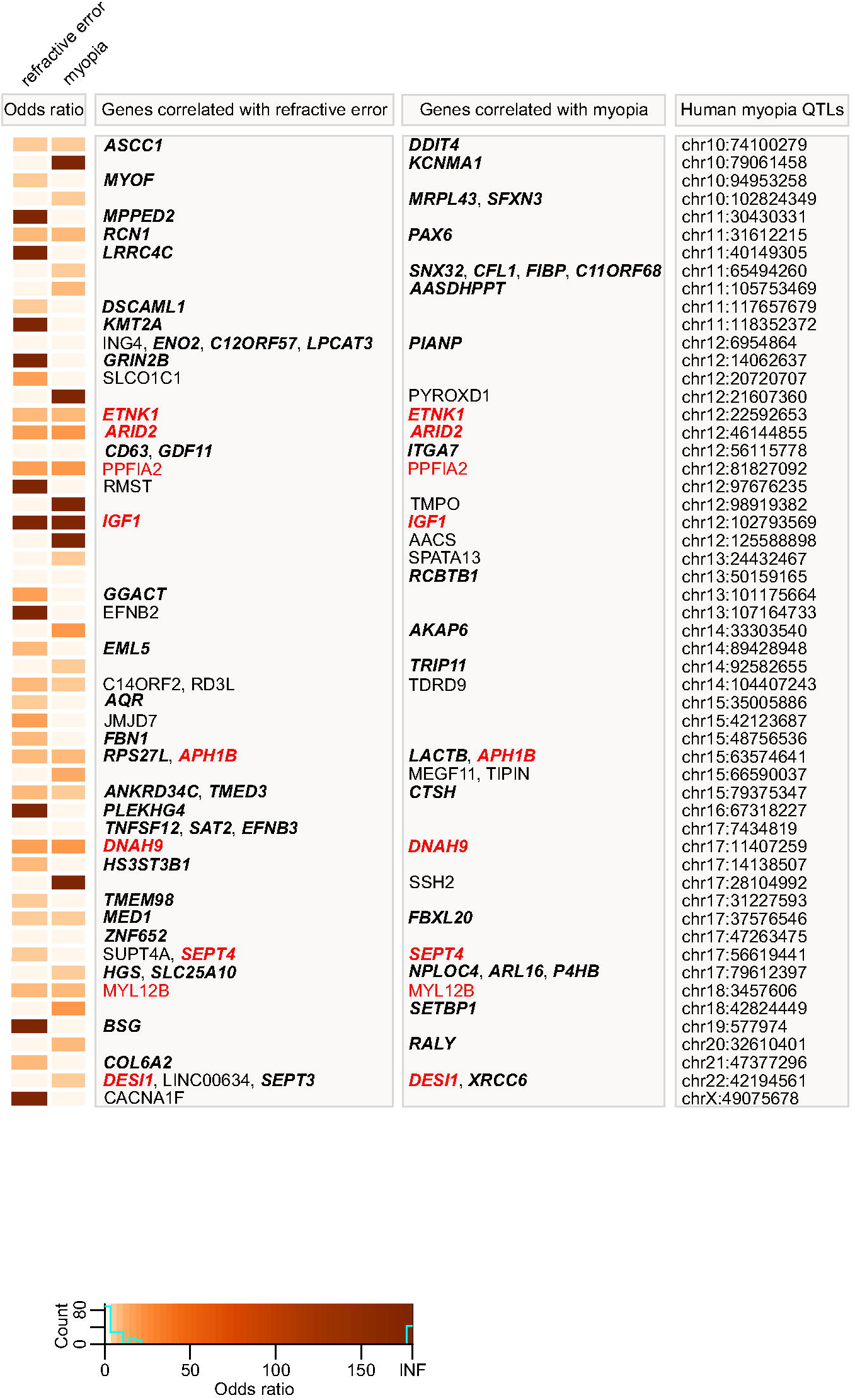
Genes underlying baseline refractive eye development and susceptibility to myopia in mice are localized in human QTLs associated with myopia (part 2). Heatmap depicting genes and odds ratios for the overlaps between 109 human myopia QTLs and genes whose expression correlates with either baseline refractive errors, susceptibility to form deprivation myopia, or both in mice. Colors indicate odds ratios. Bold italic identifies genes found to be associated with refractive error in UK Biobank, CREAM, or both human samples by the gene-based genome-wide association analysis. Red identifies genes exhibiting correlation with both baseline refractive development and susceptibility to myopia in mice.

We next analyzed biological processes linked to the genes associated with refractive development in mice and localized within human myopia QTLs (Additional file 17: Table S17, Additional file 18: Table S18, Fig 9). Surprisingly, we found that although several biological processes, such as cell growth and proliferation, circadian regulation of gene expression, and regulation of neuron differentiation, were implicated in both baseline refractive development and regulation of susceptibility to myopia, many biological processes underlying baseline refractive development and susceptibility to myopia appeared to be different. Our data on the genes associated with baseline refractive development in mice and localized in human QTLs implicate several unique processes, including camera-type eye development, regulation of ion/calcium transport, regulation of membrane polarization during action potential and long-term synaptic potentiation, nitric oxide signaling, ephrin receptor signaling, glucocorticoid receptor signaling, regulation of glutamate secretion, and regulation of JAK-STAT and MAPK signaling cascades (Additional file 17: Table S17, Fig 9A). Conversely, the genes associated with susceptibility to myopia in mice and localized in human QTLs highlighted several different processes, including regulation of cell shape and cell migration, dorsal/ventral pattern formation, vasculature morphogenesis, photoreceptor cell function and development, cellular response to DNA damage, and small-GTPase-mediated signal transduction (Additional file 18: Table S18, Fig 9B). Collectively, these data suggest that there is a significant functional overlap between genes we found to be correlated with refractive eye development in mice and genes causing myopia in humans. Moreover, our data suggest that development of refractive errors in humans is associated with genes that, in mice, regulate both baseline refractive development and susceptibility to refractive changes induced by the visual environment.

**Fig 9.**
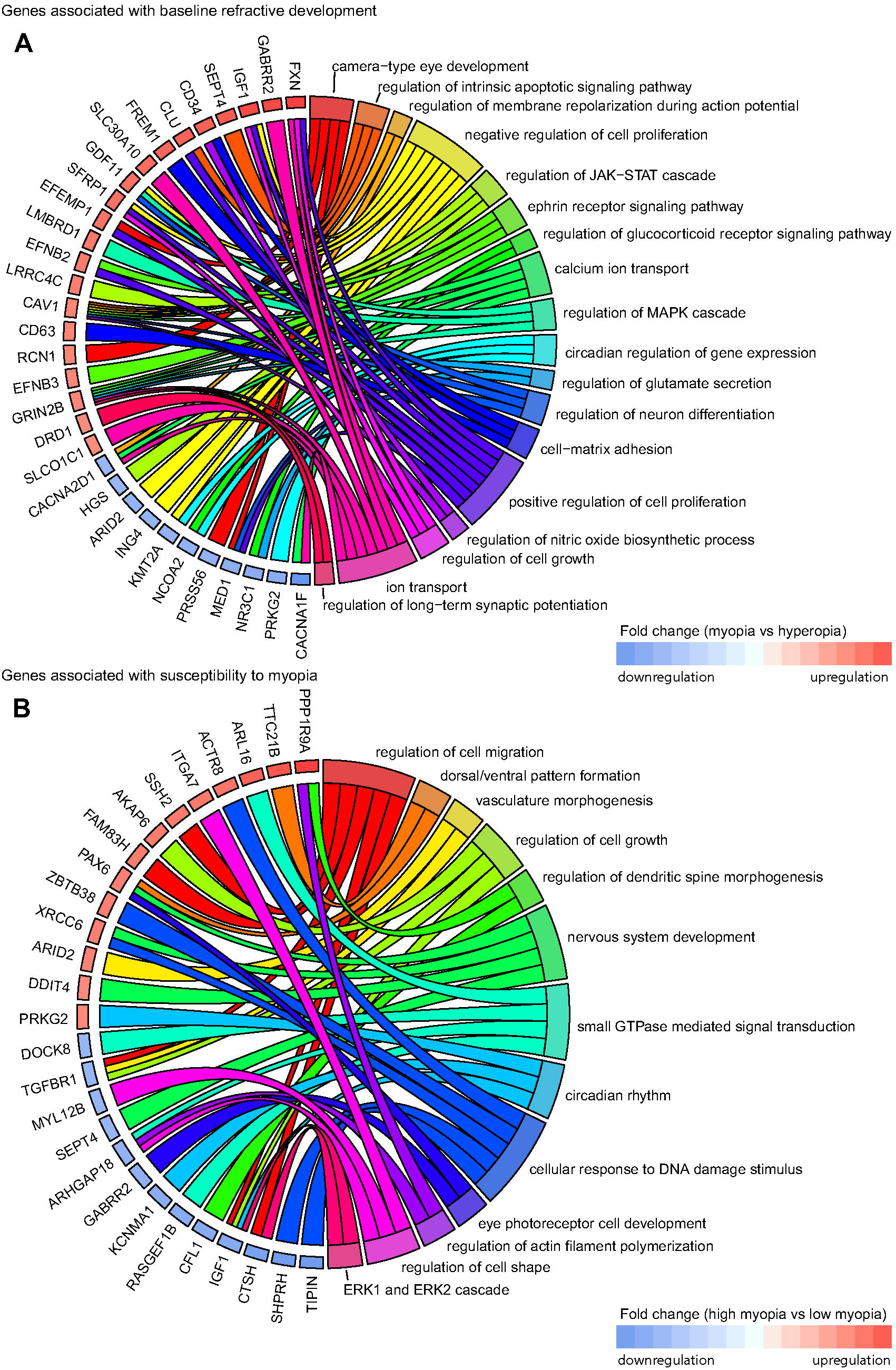
Top biological processes associated with genes linked to refractive eye development in mice and localized in known human myopia QTLs. **a** Chord diagram showing key genes (left semicircle) and top biological processes (right semicircle) associated with genes correlated with baseline refractive development in mice and localized in known human QTLs linked to myopia. **b** Chord diagram showing key genes (left semicircle) and top biological processes (right semicircle) for genes correlated with susceptibility to myopia in mice and localized in known human QTLs linked to myopia. Colored bars underneath gene names show up- or down-regulation of corresponding genes in either myopic mice versus hyperopic mice (A), or mice with high susceptibility to myopia versus mice with low susceptibility to myopia (B).

### Gene-based genome-wide association analysis of mouse genes identifies novel gene candidates for human myopia

To build on the significant overlap observed above between genes associated with refractive eye development in mice and human myopia QTLs, we examined whether the mouse genes whose expression level correlated with refractive error or myopia susceptibility were enriched for genetic variants associated with refractive error in humans. Human genes enriched for variants associated with refractive error were identified with MAGMA (Multi-marker Analysis of GenoMic Annotation), using single-marker summary statistics from a genome-wide association study (GWAS) for refractive error and age-of-onset-of-spectacle-wear reported by the CREAM Consortium (N=160,420 participants; [54]) or a GWAS for refractive error by the UK Biobank Eye & Vision Consortium (N= 88,005 participants; [71]) as input. Flanking regions of 200 kb upstream and downstream of transcription start and stop sites were included, in order to capture regulatory variants influencing the expression of nearby genes.

Of the 2,302 genes associated with baseline refractive development in mice for which there were human homologs, 277 genes in the CREAM dataset were associated with refractive error in humans (*FDR* < 0.05), with 86 genes surviving Bonferroni correction (*P_Bonferroni_*< 0.05) (Additional file 19: Table S19). Sixty-seven of these genes (including 43 genes with *P_Bonferroni_*< 0.05) were localized within previously identified human QTLs (Fig 7 and 8), while 210 of the 277 genes (including 43 genes with *P_Bonferroni_*< 0.05) have not previously been implicated in the development of refractive errors in the human population (Additional file 19: Table S19). When the above analysis was repeated using the (independent) UK Biobank dataset, 560 genes were associated with refractive error (*FDR* < 0.05), including 156 genes with genome-wide significance (*P_Bonferroni_* < 0.05) (Additional file 20: Table S20). 488 of the 560 genes (including 104 genes with *P_Bonferroni_* < 0.05) have not previously been implicated in the development of refractive errors in humans (Additional file 20: Table S20). Importantly, 190 genes associated with baseline refractive development in mice were replicated in both the CREAM and UK Biobank cohorts, including 69 genes which achieved genome-wide significance (*P_Bonferroni_*< 0.05) (Additional file 23: Table S23).

Of the 1,917 genes associated with the regulation of susceptibility to myopia in mice, gene-based analysis using the CREAM dataset identified 223 genes associated with refractive error in humans (*FDR* < 0.05), including 72 genes at genome-wide significance (*P_Bonferroni_*< 0.05) (Additional file 21: Table S21). Forty-six of these genes (including 33 genes with *P_Bonferroni_* < 0.05) were localized within previously identified human QTLs (Fig 7 and 8), whereas 177 of the 223 genes (including 39 genes with *P_Bonferroni_*< 0.05) have not previously been implicated in the development of refractive errors in humans (Additional file 21: Table S21). MAGMA analysis of the 1,917 genes using the UK Biobank dataset revealed that 465 genes were associated with refractive error (*FDR* < 0.05), including 119 genes that withstood Bonferroni correction (*P_Bonferroni_* < 0.05) (Additional file 22: Table S22). Forty-nine of these genes (including 35 genes with *P_Bonferroni_*< 0.05) were localize within known human myopia QTLs (Fig 7 and 8), whereas 416 of the 465 genes (including 84 genes with *P_Bonferroni_*< 0.05) were not previously known to be associated with the development of refractive errors in the human population (Additional file 22: Table S22). One hundred and fifty-two genes involved in the regulation of susceptibility to myopia in mice were found to be linked to the development of refractive errors in both CREAM and UK Biobank cohorts, including 53 genes which reached genome-wide significance (*P_Bonferroni_* < 0.05) in both samples (Additional file 24: Table S24).

Analysis of the biological processes associated with the above genes linked to either baseline refractive development or regulation of susceptibility to myopia in humans (Additional file 25: Table S25, Additional file 26: Table S26, Fig 10A and 11A) revealed processes primarily related to calcium-mediated cell adhesion, synapse assembly, synaptic transmission, protein translation, small GTPase-mediated signal transduction, and GABA receptor signaling. Several canonical signaling pathways were also identified, including EIF2 and mTOR signaling, eIF4 and p70S6K signaling, epithelial adherence junction signaling, sumoylation pathway, and regulation of cellular mechanics by calpain protease, among others (Additional file 27: Table S27, Additional file 28: Table S28, Fig 10B and 11B).

**Fig 10.**
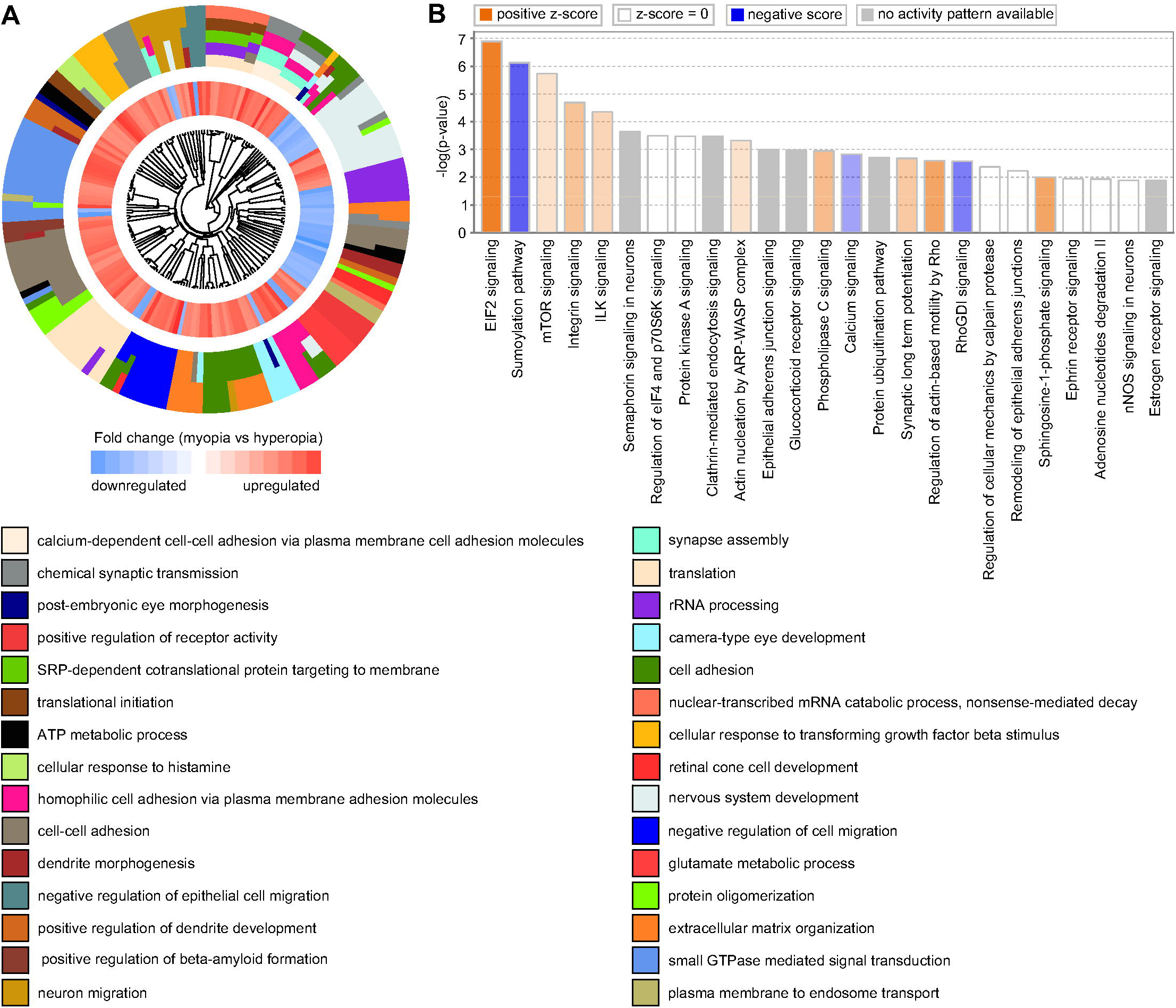
Summary of key biological processes and pathways affected by genes underlying refractive eye development and found to be linked to myopia in UK Biobank and CREAM human cohorts. **a** Hierarchical clustering diagram showing top 30 biological processes affected by genes underlying baseline refractive eye development in humans. Outer circle shows hierarchical clusters of biological processes (identified by different colors) linked to baseline refractive development; inner circle shows clusters of the corresponding genes up- or down-regulated in myopic mice versus hyperopic mice. **b** Top twenty-five canonical pathways affected by the genes underlying baseline refractive eye development in humans. Horizontal yellow line indicates *P* = 0.05. Z-score shows activation or suppression of the corresponding pathways.

**Fig 11.**
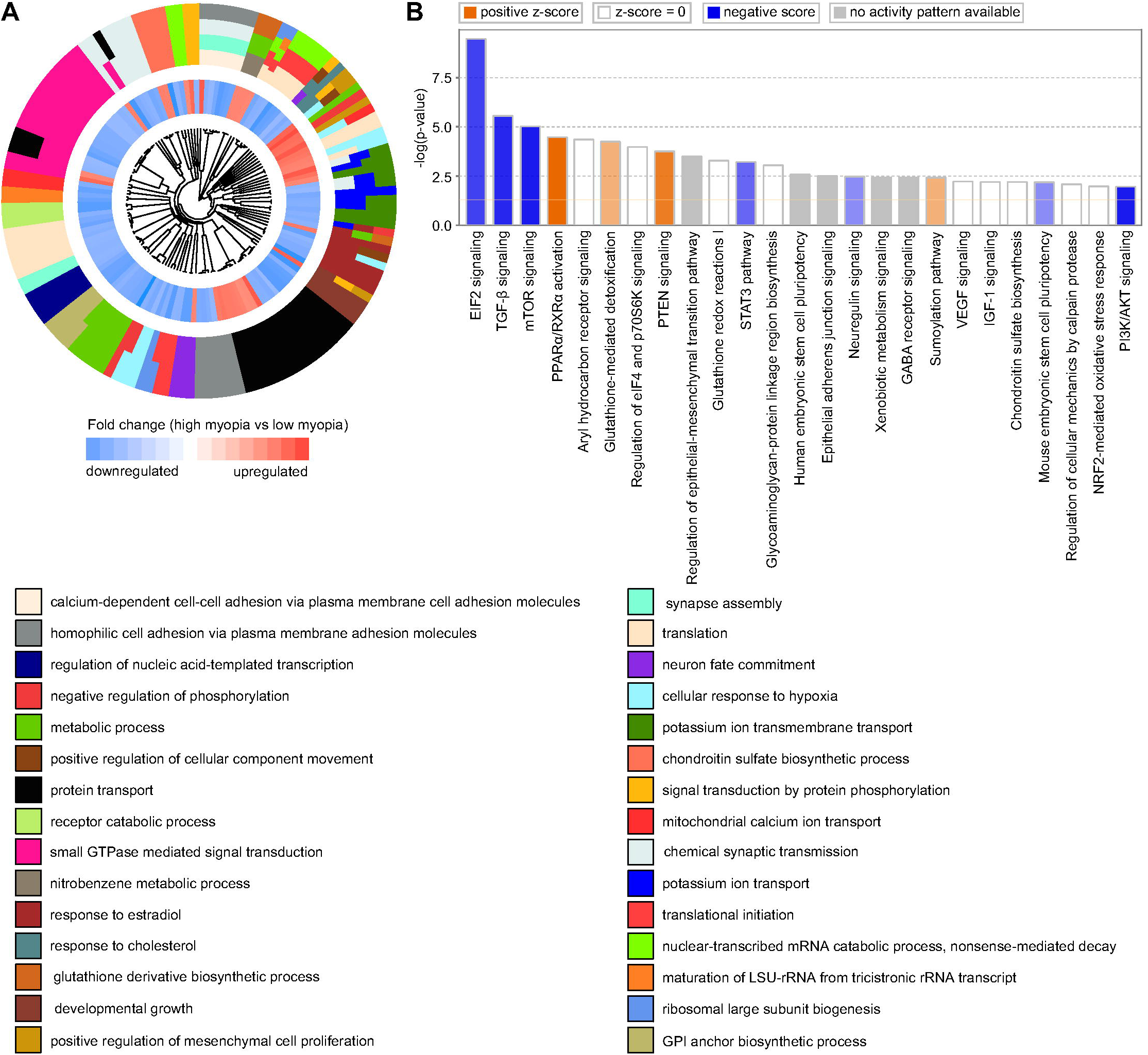
Summary of key biological processes and pathways affected by genes regulating susceptibility to myopia and found to be linked to myopia in UK Biobank and CREAM human cohorts. **a** Hierarchical clustering diagram showing top 30 biological processes affected by genes regulating susceptibility to myopia in humans. Outer circle shows hierarchical clusters of biological processes (identified by different colors) underlying regulation of susceptibility to myopia; inner circle shows clusters of the corresponding genes up- or down-regulated in mice with high susceptibility to myopia versus mice with low susceptibility to myopia. **b** Top twenty-five canonical pathways affected by the genes regulating susceptibility to myopia in humans. Horizontal yellow line indicates *P* = 0.05. Z-score shows activation or suppression of the corresponding pathways.

However, there were also substantial differences between the two sets of human genes identified using either the mouse baseline refractive error or the mouse myopia susceptibility gene sets. The mouse baseline refractive error-derived human gene set was associated with nervous system development, post-embryonic camera-type eye development, retinal cone development, neuron migration, dendrite morphogenesis, regulation of glutamate metabolism, extracellular matrix organization, and beta-amyloid formation (Additional file 25: Table S25, Fig 10A). This gene set was associated with several canonical pathways distinct from those identified using the mouse susceptibility to myopia-derived human gene set. These pathways included integrin signaling, semaphorin signaling in neurons, glucocorticoid receptor signaling, phospholipase C signaling, synaptic long term potentiation, ephrin receptor signaling, nNOS signaling in neurons, and estrogen receptor signaling (Additional file 27: Table S27, Additional file 28: Table S28, Fig 10B and 11B).

The mouse susceptibility to myopia-derived human gene set, on the other hand, was associated with biological processes related to developmental growth, neuron fate commitment, regulation of mesenchymal cell proliferation, potassium ion transmembrane transport, protein transport, response to estradiol and cholesterol, as well as cellular response to hypoxia (Additional file 26: Table S26, Fig 11A). Analysis of canonical pathways suggested that pathways associated with TGF-β signaling, PPARα/RXRα activation, PTEN signaling, regulation of the epithelial-mesenchymal transition, STAT3 signaling, regulation of stem cell pluripotency, VEGF and IGF-1 signaling, NRF2-mediated oxidative stress response, and PI3K/AKT signaling, among others, were involved in the regulation of susceptibility to myopia (Additional file 28: Table S28, Fig 11B).

## Discussion

Human population studies and studies in animal models strongly suggest that both environmental and genetic factors play important roles in refractive eye development. Numerous linkage and genome-wide association studies in humans identified over 270 chromosomal loci linked to the development of myopia in humans [74]; however, very little is known how genetic variation causing myopia affects gene expression and how changes in gene expression associated with differences in genetic backgrounds affect refractive eye development and susceptibility to myopia.

Our data suggest that differences in genetic backgrounds play a very important role in refractive eye development and regulation of susceptibility to myopia. Moreover, we found that variations in genetic background produce a continuous distribution of refractive errors and susceptibilities to myopia in a mouse population, characteristic of quantitative traits. Genetic variations in different mouse strains also produce unique patterns of gene expression, which strongly correlate with either baseline refractive errors or susceptibility to induced myopia. Surprisingly, we found that the baseline refractive development and susceptibility to myopia are controlled by largely distinct sets of genes, which suggests that signaling pathways that regulate the trajectory of refractive eye development towards emmetropia, myopia, or hyperopia might be different from the pathways that modulate the impact of optical defocus and other environmental factors on refractive development. Nevertheless, expression of 714 genes correlated with both baseline refractive errors and susceptibility to myopia, suggesting that at least some genes control both the pathways regulating the trajectory of refractive development and the impact of visual input on it.

Genes that influenced either baseline refractive development or susceptibility to myopia affected a long list of biological and molecular functions; however, most noteworthy is the finding that over 29 canonical signaling pathways were involved in both baseline refractive development and regulation of susceptibility to myopia. Interestingly, the majority of these pathways were suppressed in animals with high susceptibility to myopia and activated in animals with negative baseline refractive errors. The exceptions from this rule were sumoylation, RhoGDI signaling, PTEN, and HIPPO signaling pathways, which we found to be activated in animals with high susceptibility to myopia and suppressed in animals with negative baseline refractive errors. The observation that the same pathways were effected in the opposite directions in animals with high susceptibility to myopia and animals with highly negative baseline refractive errors may be explained by the role of optical defocus in the development of myopia. Considering that mice are housed in small cages and the exposure to distant vision is limited, animals with hyperopic refractive errors would be exposed to high levels of hyperopic optical defocus (which was shown to cause myopia) compared to the animals with negative refractive errors; thus explaining why signaling pathways are effected in the same direction in animals with high susceptibility to myopia and animals with hyperopic baseline refractive errors.

Remarkably, we also found that many genes whose expression correlated with either baseline refractive errors or susceptibility to myopia in mice were localized in the known human QTLs linked to myopia. The majority of these genes (90 genes) were exclusively involved in baseline refractive eye development, 51 genes were linked to the regulation of susceptibility to myopia, and 23 genes affected both baseline refractive development and susceptibility to myopia.

Gene-based genome-wide association analysis of the genes we found in mice against CREAM and UK Biobank human samples revealed that 647 genes whose expression correlated with baseline refractive errors in mice were also associated with refractive errors in humans, including 173 genes that withstood Bonferroni correction. Using gene-based analysis, we also found that 536 genes whose expression correlated with susceptibility to myopia were associated with refractive errors in humans, including 138 genes which exhibited genome-wide significance (*P_Bonferroni_* < 0.05). One hundred and ninety-eight of these genes were involved in both baseline refractive development and regulation of susceptibility to myopia, including 34 genes which withstood Bonferroni correction. Although many of these genes were previously implicated in the development of human myopia, 572 genes (92 genes with *P_Bonferroni_* < 0.05) whose expression correlated with baseline refractive development and 486 genes (79 genes with *P_Bonferroni_*< 0.05) whose expression correlated with susceptibility to myopia were linked to human myopia for the first time, including 211 genes (54 genes with *P_Bonferroni_*< 0.05) whose expression correlated with both baseline refractive development and susceptibility to myopia.

Genes that we found to be involved in refractive error development in mice and humans affect a multitude of biological functions in the retina; however, many biological functions underlying refractive development appear to be highly conserved across species. These include camera-type eye development and post-embryonic eye morphogenesis, ephrin receptor signaling, glucocorticoid receptor signaling, regulation of circadian rhythms and circadian regulation of gene expression, glutamate signaling, regulation of neurogenesis and dendrite morphogenesis, regulation of nitric oxide biosynthesis, regulation of long-term synaptic potentiation, synapse assembly and chemical synapse transmission, calcium-dependent signaling, regulation of translation, small GTPase mediated signal transduction, photoreceptor function and development, cell-cell adhesion, regulation of beta-amyloid formation, regulation of mesenchymal cell proliferation, cellular response to hypoxia. We also found that many retinal signaling pathways involved in refractive development in mice are also subjected to genetic variation causing myopia in humans. These include EIF2 signaling, regulation of eIF4 and p70S6K signaling, mTOR signaling, integrin signaling, semaphorin signaling in neurons, protein kinase A signaling, clathrin-mediated endocytosis signaling, epithelial adherens junction signaling, synaptic long term potentiation, ephrin receptor signaling, nNOS and eNOS signaling, estrogen receptor signaling, PPARα/RXRα activation, signaling by Rho family GTPases, Wnt/Ca+ pathway, dopamine-DARPP32 feedback signaling, PTEN signaling, IGF-1 signaling, insulin receptor signaling, NF-κB signaling, RAR signaling, amyloid processing, among other pathways.

Interestingly, 27 genes that we found to be associated with refractive error development in mice were also among the genes differentially expressed in the retina of green monkeys with form-deprivation-induced myopia [39], and 233 genes were among the genes differentially expressed in the retina of marmosets exposed to positive and negative optical defocus [75]. There was also a 47-gene overlap with the genes found by Riddell et al. to be differentially expressed in the retina of chicks exposed to optical defocus [76], while 292 genes found in mice overlapped with genes found to be differentially expressed in the retina of chicks with lens-induced myopia by Stone et al. [77]. Many of the genes that we found to be involved in refractive eye development in this study had been previously implicated in various physiological and pathological processes in the retina. For example, mutations in *TTC21B* gene, which we found to be involved in the regulation of susceptibility to myopia, were shown to cause a syndromic form of retinal dystrophy [78]. Mutations in *EFEMP1* (gene involved in baseline refractive development) were found to be associated with Malattia Leventinese retinal dystrophy characterized by the presence of RPE deposits [79]. Several studies linked a manganese transporter *SLC30A10*, which was found in this study to be involved in baseline refractive development, to parkinsonism, thus implicating this gene in the regulation of dopamine signaling [80, 81]. Ephrin receptor A10 (*EPHA10*), which we found to be involved in the regulation of susceptibility to myopia, was shown to influence cone photoreceptor morphogenesis, implicating signaling at the level of photoreceptors in refractive eye development [82]. A GTP-binding protein GNL2, involved in baseline refractive development, was demonstrated to play important role in retinal neurogenesis [83]. Finally, *AGRN* gene, found by this study to be involved in baseline refractive development, was shown to interact with EGR1 (previously implicated in refractive eye development) and regulate synaptic physiology in the retina [84, 85].

Although many pathways that we found to underlie refractive development in mice and humans in this study are novel, many of them are conserved across vertebrate species (Table 1). Seven out of 47 pathways found to be involved in optical defocus response in chickens by Riddell et al. [62] were also found to be involved in refractive eye development in this study, including PPAR signaling, tight junction signaling, Huntington’s disease signaling, cysteine metabolism, actin cytoskeleton signaling, TGF-β signaling, and Glutathione metabolism (Additional file 5: Table S5, Additional file 8: Table S8, Additional file 27: Table S27, Additional file 28: Table S28). We replicated 12 out of 20 canonical pathways identified by Stone et al. [77] in chickens with lens-induced myopia, including glutamate receptor signaling, β-adrenergic signaling, synaptic long term depression, synaptic long term potentiation, amyotrophic lateral sclerosis signaling, CREB signaling in neurons, relaxin signaling, purine metabolism, CDK5 signaling, protein kinase A signaling, hypoxia signaling, and tight junction signaling (Additional file 5: Table S5, Additional file 8: Table S8, Additional file 27: Table S27, Additional file 28: Table S28). We also found that 35 out of 75 pathways that we recently found to be involved in the development of hyperopia and myopia in marmosets [75] were also among the pathways which we found to be involved in refractive eye development in this study, including Ephrin Receptor Signaling, β-adrenergic Signaling, Protein Kinase A Signaling, Relaxin Signaling, Androgen Signaling, Dopamine-DARPP32 Feedback Signaling, nNOS Signaling, RAN Signaling, HIPPO signaling, Wnt/Ca+ pathway, PTEN Signaling, Synaptic Long Term Potentiation, Gap Junction Signaling, Regulation of eIF4 and p70S6K Signaling, mTOR Signaling, Glucocorticoid Receptor Signaling, α-Adrenergic Signaling, and Epithelial Adherens Junction Signaling, among others (Additional file 5: Table S5, Additional file 8: Table S8, Additional file 27: Table S27, Additional file 28: Table S28). An important role of amyloid signaling pathway, which we found to be involved in the regulation of susceptibility to myopia (Additional file 8: Table S8), is an agreement with our recent finding that a component of the amyloid signaling pathway APLP2 regulates susceptibility to myopia in mice and humans [37]. Our data also suggest that the phototransduction pathway is involved in refractive eye development (Additional file 5: Table S5), in agreement with a recent GWAS study [54] and recent marmoset data [75].

**Table 1.**
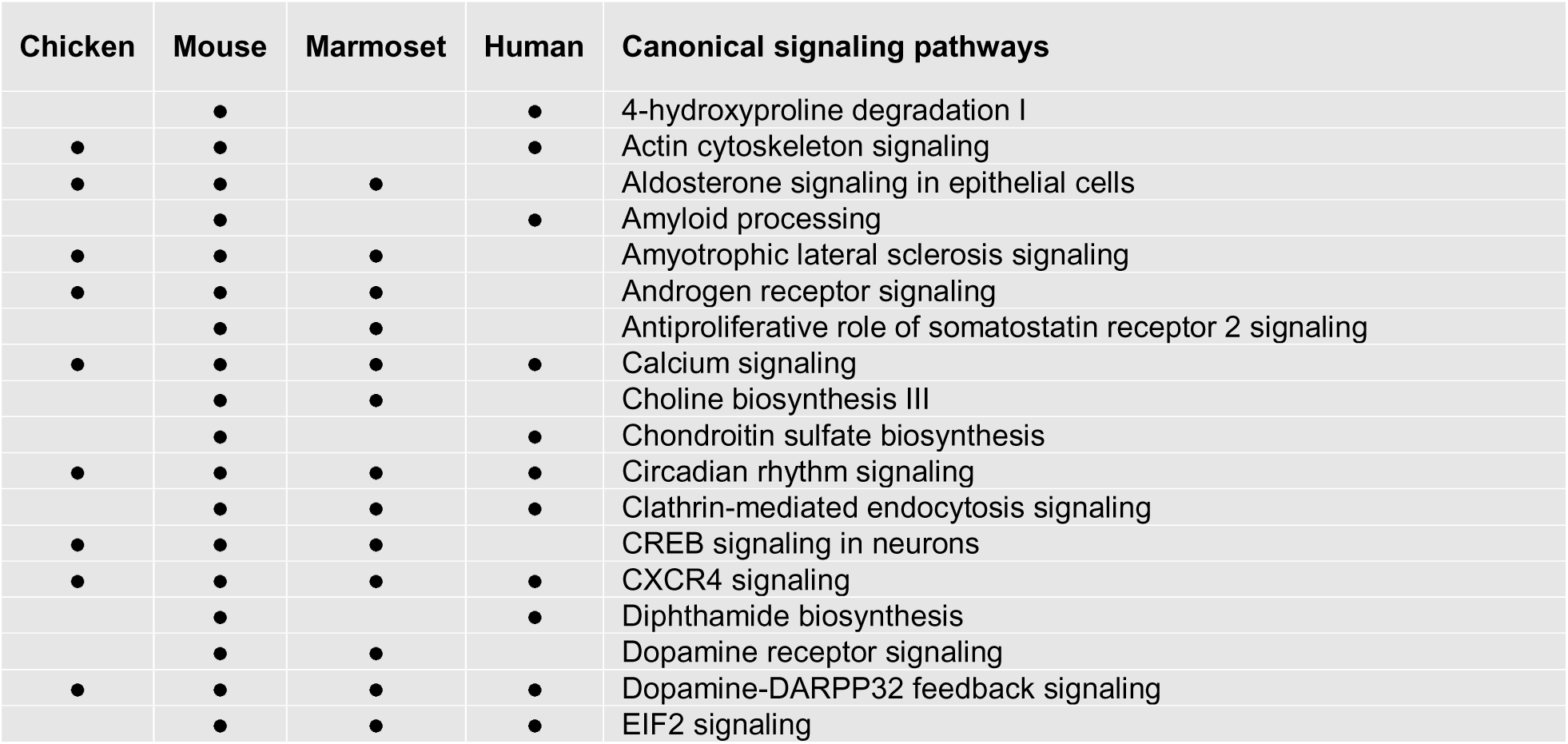
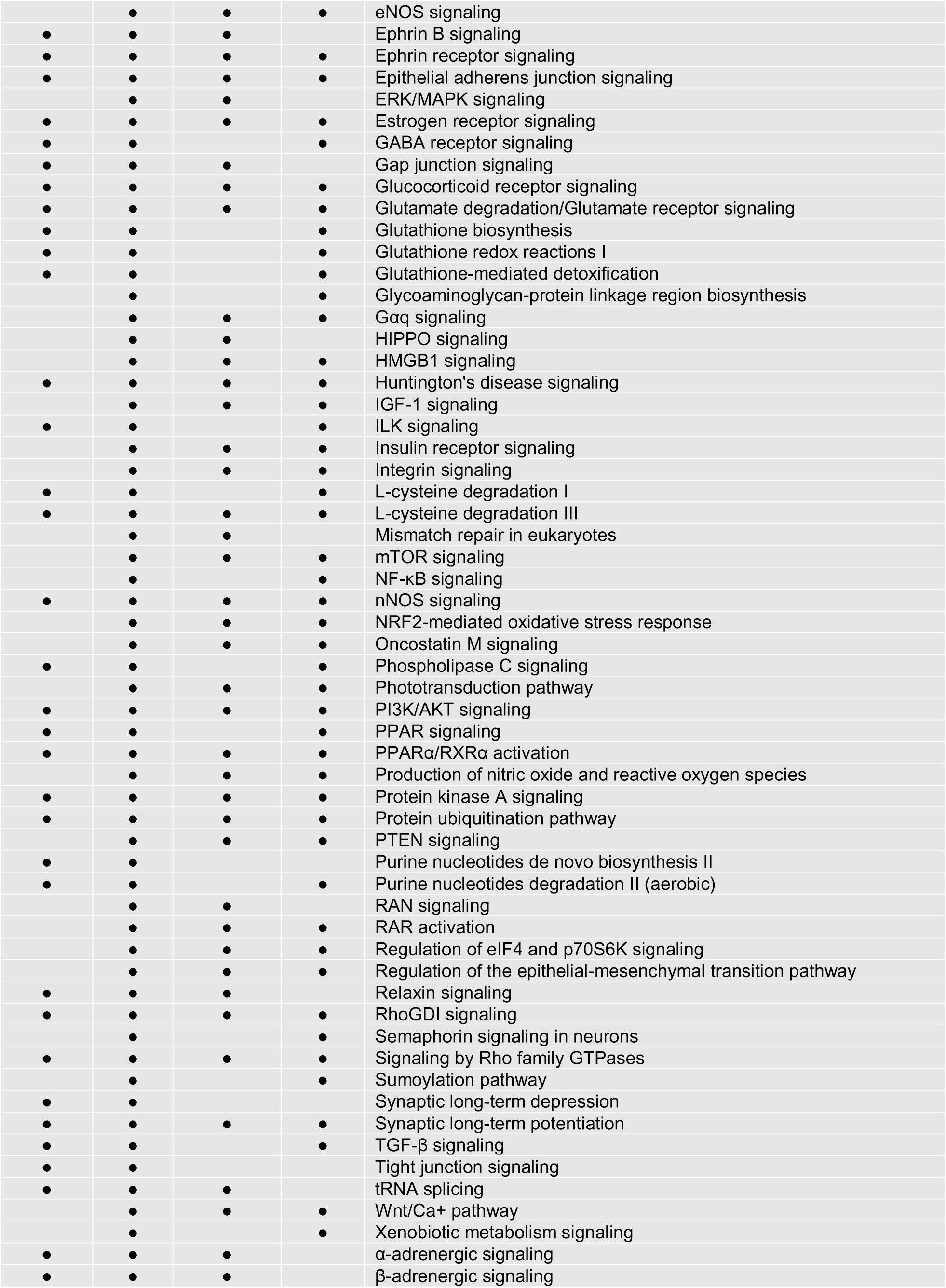
Evolutionary conservation of retinal signaling pathways involved in refractive eye development.

## Conclusions

We have identified 2,302 genes which are involved in baseline refractive eye development and 1,917 genes which regulate susceptibility to myopia in mice. Our data suggest that at least 985 of these genes are subjected to genetic variation in the human population and are involved in refractive error development in humans. Eight hundred forty-seven of these genes were implicated in the development of human myopia for the first time. A large number of common genes and canonical pathways that we found to be involved in the regulation of refractive eye development in mice, chickens and humans suggests strong evolutionary conservation of the signaling pathways underlying refractive error development. It appears that the number of genes involved in the regulation of refractive eye development may be as high as 3,505 in the retina alone, suggesting that refractive eye development maybe regulated by hundreds to thousands of genes across ocular tissues. Interestingly, in spite of significant overlap between genes that control baseline refractive development and genes regulating susceptibility to myopia, these two processes appear to be controlled by largely distinct sets of genes. The genes that we found to be involved in refractive eye development control, however, a well-defined set of retinal signaling pathways that we begin to consistently find to be involved in refractive eye development across different species. This provides a solid framework for future studies and for the development of anti-myopia drugs.

## Acknowledgements

We are grateful to the CREAM Consortium, the UK Biobank Eye and Vision Consortium, and all the individuals who took part in the CREAM and UK Biobank studies, the personnel who recruited them, interviewers, computer and laboratory technicians, clerical workers, research scientists, volunteers, managers, receptionists and nurses. This research has been conducted using the CREAM datasets and the UK Biobank Resource (application #17351). Special thanks to Jeremy Guggenheim who provided valuable feedback on an early version of the manuscript.

## Additional files

**Additional file 1: Table S1.** Baseline refractive errors in Collaborative Cross mice measured at P40 (diopters).

**Additional file 2: Table S2.** Form-deprivation myopia in Collaborative Cross mice (deprived eye versus control eye, diopters).

**Additional file 3: Table S3.** List of genes whose expression correlates with baseline refractive error in Collaborative Cross mice.

**Additional file 4: Table S4.** Gene ontology categories significantly associated with genes whose expression correlates with baseline refractive error in Collaborative Cross mice (BP, biological process; CC, cellular component; MF, molecular function).

**Additional file 5: Table S5.** Canonical signaling pathways affected by genes whose expression correlates with baseline refractive error in Collaborative Cross mice.

**Additional file 6: Table S6.** List of genes whose expression correlates with susceptibility to myopia in Collaborative Cross mice.

**Additional file 7: Table S7.** Gene ontology categories significantly associated with genes whose expression correlates with susceptibility to myopia in Collaborative Cross mice (BP, biological process; CC, cellular component; MF, molecular function).

**Additional file 8: Table S8.** Canonical signaling pathways affected by genes whose expression correlates with susceptibility to myopia in Collaborative Cross mice.

**Additional file 9: Table S9.** List of genes whose expression correlates with both baseline refractive error and susceptibility to myopia in Collaborative Cross mice.

**Additional file 10: Table S10.** Gene ontology categories significantly associated with genes whose expression correlates with both baseline refractive error and susceptibility to myopia in Collaborative Cross mice (BP, biological process; CC, cellular component; MF, molecular function).

**Additional file 11: Table S11.** Canonical signaling pathways affected by genes whose expression correlates with both baseline refractive error and susceptibility to myopia in Collaborative Cross mice.

**Additional file 12: Table S12.** List of human myopia QTLs and candidate genes located within 200 kb of the lead SNP (or within critical region).

**Additional file 13: Table S13.** List of genes localized within human myopia QTLs and whose expression correlates with baseline refractive error in mice.

**Additional file 14: Table S14.** List of genes localized within human myopia QTLs and whose expression correlates with susceptibility to myopia in mice.

**Additional file 15: Table S15.** List of genes localized within human myopia QTLs and whose expression correlates with both baseline refractive error and susceptibility to myopia in mice.

**Additional file 16: Table S16.** List of human myopia QTLs showing overlap with genes exhibiting correlation with either refractive error or susceptibility to myopia in mice.

**Additional file 17: Table S17.** Gene ontology categories significantly associated with genes localized within human myopia QTLs and whose expression correlates with baseline refractive error in mice (BP, biological process; CC, cellular component; MF, molecular function).

**Additional file 18: Table S18.** Gene ontology categories significantly associated with genes localized within human myopia QTLs and whose expression correlates with susceptibility to myopia in mice (BP, biological process; CC, cellular component; MF, molecular function).

**Additional file 19: Table S19.** List of mouse genes displaying significant association with refractive error in CREAM cohort and whose expression correlates with baseline refractive error in mice (Blue identifies P_Bonferroni_ < 0.05 in both cohorts).

**Additional file 20: Table S20.** List of mouse genes displaying significant association with refractive error in UK Biobank cohort and whose expression correlates with baseline refractive error in mice (Blue identifies P_Bonferroni_ < 0.05 in both cohorts).

**Additional file 21: Table S21.** List of mouse genes displaying significant association with refractive error in CREAM cohort and whose expression correlates with susceptibility to myopia in mice (Blue identifies P_Bonferroni_ < 0.05 in both cohorts).

**Additional file 22: Table S22.** List of mouse genes displaying significant association with refractive error in UK Biobank cohort and whose expression correlates with susceptibility to myopia in mice (Blue identifies P_Bonferroni_ < 0.05 in both cohorts).

**Additional file 23: Table S23.** List of mouse genes displaying significant association with refractive error in both UK Biobank and CREAM cohorts, and whose expression correlates with baseline refractive error in mice (Blue identifies P_Bonferroni_ < 0.05 in both cohorts).

**Additional file 24: Table S24.** List of mouse genes displaying significant association with refractive error in both UK Biobank and CREAM cohorts, and whose expression correlates with susceptibility to myopia in mice (Blue identifies P_Bonferroni_ < 0.05 in both cohorts).

**Additional file 25: Table S25.** Biological processes significantly associated with genes linked to refractive error in UK Biobank and CREAM cohorts, and whose expression correlates with baseline refractive error in mice (BP, biological process).

**Additional file 26: Table S26.** Biological processes significantly associated with genes linked to refractive error in UK Biobank and CREAM cohorts, and whose expression correlates with susceptibility to myopia in mice (BP, biological process).

**Additional file 27: Table S27.** Canonical signaling pathways significantly associated with genes linked to refractive error in UK Biobank and CREAM cohorts, and whose expression correlates with baseline refractive error in mice.

**Additional file 28: Table S28.** Canonical signaling pathways significantly associated with genes linked to refractive error in UK Biobank and CREAM cohorts, and whose expression correlates with susceptibility to myopia in mice.

## References

1. Pararajasegaram R. VISION 2020-the right to sight: from strategies to action. Am J Ophthalmol. 1999;128(3):359–60.

2. Kempen JH, Mitchell P, Lee KE, Tielsch JM, Broman AT, Taylor HR, et al. The prevalence of refractive errors among adults in the United States, Western Europe, and Australia. Arch Ophthalmol. 2004;122(4):495–505.

3. Javitt JC, Chiang YP. The socioeconomic aspects of laser refractive surgery. Arch Ophthalmol. 1994;112(12):1526–30.

4. Vitale S, Sperduto RD, Ferris FL, 3rd. Increased prevalence of myopia in the United States between 1971-1972 and 1999-2004. Arch Ophthalmol. 2009;127(12):1632–9.

5. Holden BA, Fricke TR, Wilson DA, Jong M, Naidoo KS, Sankaridurg P, et al. Global Prevalence of Myopia and High Myopia and Temporal Trends from 2000 through 2050. Ophthalmology. 2016;123(5):1036–42.

6. Lin LL, Shih YF, Hsiao CK, Chen CJ. Prevalence of myopia in Taiwanese schoolchildren: 1983 to 2000. Ann Acad Med Singapore. 2004;33(1):27–33.

7. Lam CS, Goldschmidt E, Edwards MH. Prevalence of myopia in local and international schools in Hong Kong. Optom Vis Sci. 2004;81(5):317–22.

8. Alexander LJ. Primary care of the posterior segment. 2 ed. Connecticut: Appleton & Lange; 1994.

9. Saw SM, Gazzard G, Shih-Yen EC, Chua WH. Myopia and associated pathological complications. Ophthalmic Physiol Opt. 2005;25(5):381–91.

10. Flitcroft DI. The complex interactions of retinal, optical and environmental factors in myopia aetiology. Prog Retin Eye Res. 2012;31(6):622–60.

11. Hurst J, Johnson D, Law C, Schweitzer K, Sharma S. Value of subjective visual reduction in patients with acute-onset floaters and/or flashes. Can J Ophthalmol. 2015;50(4):265–8.

12. Shunmugam M, Ang GS, Lois N. Giant retinal tears. Surv Ophthalmol. 2014;59(2):192–216.

13. Rumelt S, Sarrazin L, Averbukh E, Halpert M, Hemo I. Paediatric vs adult retinal detachment. Eye (Lond). 2007;21(12):1473–8.

14. Brasil OF, Brasil MV, Japiassu RM, Biancardi AL, Souza DD, Oliveira RC, et al. [Fundus changes evaluation in degenerative myopia]. Arquivos brasileiros de oftalmologia. 2006;69(2):203–6.

15. Ndiaye PA, Koffane RR, Wade A, Ndiaye CS, Gomez JC, Ndiaye MR. [Frequency of retinal changes in myopia in a black population]. J Fr Ophtalmol. 2001;24(9):927–9.

16. Bonnet M. [Myopia and rhegmatogenous retinal detachment]. Rev Prat. 1993;43(14):1779–83.

17. Pierro L, Camesasca FI, Mischi M, Brancato R. Peripheral retinal changes and axial myopia. Retina. 1992;12(1):12–7.

18. Grossniklaus HE, Green WR. Pathologic findings in pathologic myopia. Retina. 1992;12(2):127–33.

19. Burton TC. The influence of refractive error and lattice degeneration on the incidence of retinal detachment. Trans Am Ophthalmol Soc. 1989;87:143–55; discussion 55-7.

20. Pruett RC, Weiter JJ, Goldstein RB. Myopic cracks, angioid streaks, and traumatic tears in Bruch’s membrane. Am J Ophthalmol. 1987;103(4):537–43.

21. Chaine G, Sebag J, Coscas G. The induction of retinal detachment. Trans Ophthalmol Soc U K. 1983;103 (Pt 4):480–5.

22. Menezo JL, Suarez-Reynolds R, Frances J, Vila E. Shape, number and localization of retinal tears in myopic over 8D, aphakic and traumatic cases of retinal detachment. An experience report. Ophthalmologica. 1977;175(1):10–8.

23. Kanski JJ. Giant retinal tears. Am J Ophthalmol. 1975;79(5):846–52.

24. Verhoeven VJ, Wong KT, Buitendijk GH, Hofman A, Vingerling JR, Klaver CC. Visual consequences of refractive errors in the general population. Ophthalmology. 2015;122(1):101–9.

25. Qiu M, Wang SY, Singh K, Lin SC. Association between myopia and glaucoma in the United States population. Invest Ophthalmol Vis Sci. 2013;54(1):830–5.

26. Praveen MR, Vasavada AR, Jani UD, Trivedi RH, Choudhary PK. Prevalence of cataract type in relation to axial length in subjects with high myopia and emmetropia in an Indian population. Am J Ophthalmol. 2008;145(1):176–81.

27. Loyo-Berrios NI, Blustein JN. Primary-open glaucoma and myopia: a narrative review. WMJ: official publication of the State Medical Society of Wisconsin. 2007;106(2):85–9, 95.

28. Pizzarello L, Abiose A, Ffytche T, Duerksen R, Thulasiraj R, Taylor H, et al. VISION 2020: The Right to Sight: a global initiative to eliminate avoidable blindness. Arch Ophthalmol. 2004;122(4):615–20.

29. Morgan IG. The biological basis of myopic refractive error. Clin Exp Optom. 2003;86(5):276–88.

30. Young TL. Molecular genetics of human myopia: an update. Optom Vis Sci. 2009;86(1):E8–E22.

31. Baird PN, Schache M, Dirani M. The GEnes in Myopia (GEM) study in understanding the aetiology of refractive errors. Prog Retin Eye Res. 2010;29(6):520–42.

32. Wojciechowski R. Nature and nurture: the complex genetics of myopia and refractive error. Clin Genet. 2011;79(4):301–20.

33. Parssinen O, Lyyra AL. Myopia and myopic progression among schoolchildren: a three-year follow-up study. Invest Ophthalmol Vis Sci. 1993;34(9):2794–802.

34. Goss DA. Nearwork and myopia. Lancet. 2000;356(9240):1456–7.

35. Hepsen IF, Evereklioglu C, Bayramlar H. The effect of reading and near-work on the development of myopia in emmetropic boys: a prospective, controlled, three-year follow-up study. Vision Res. 2001;41(19):2511–20.

36. Saw SM, Chua WH, Hong CY, Wu HM, Chan WY, Chia KS, et al. Nearwork in early-onset myopia. Invest Ophthalmol Vis Sci. 2002;43(2):332–9.

37. Tkatchenko AV, Tkatchenko TV, Guggenheim JA, Verhoeven VJ, Hysi PG, Wojciechowski R, et al. APLP2 Regulates Refractive Error and Myopia Development in Mice and Humans. PLoS Genet. 2015;11(8):e1005432.

38. Troilo D, Li T, Glasser A, Howland HC. Differences in eye growth and the response to visual deprivation in different strains of chicken. Vision Res. 1995;35(9):1211–6.

39. Tkatchenko AV, Walsh PA, Tkatchenko TV, Gustincich S, Raviola E. Form deprivation modulates retinal neurogenesis in primate experimental myopia. Proc Natl Acad Sci U S A. 2006;103(12):4681–6.

40. Schaeffel F, Burkhardt E, Howland HC, Williams RW. Measurement of refractive state and deprivation myopia in two strains of mice. Optom Vis Sci. 2004;81(2):99–110.

41. Chen YP, Hocking PM, Wang L, Povazay B, Prashar A, To CH, et al. Selective breeding for susceptibility to myopia reveals a gene-environment interaction. Invest Ophthalmol Vis Sci. 2011;52(7):4003–11.

42. Zhou G, Williams RW. Mouse models for the analysis of myopia: an analysis of variation in eye size of adult mice. Optom Vis Sci. 1999;76(6):408–18.

43. Puk O, Dalke C, Favor J, de Angelis MH, Graw J. Variations of eye size parameters among different strains of mice. Mamm Genome. 2006;17(8):851–7.

44. Wong AA, Brown RE. Visual detection, pattern discrimination and visual acuity in 14 strains of mice. Genes Brain Behav. 2006;5(5):389–403.

45. Peet JA, Cotch MF, Wojciechowski R, Bailey-Wilson JE, Stambolian D. Heritability and familial aggregation of refractive error in the Old Order Amish. Invest Ophthalmol Vis Sci. 2007;48(9):4002–6.

46. Lyhne N, Sjolie AK, Kyvik KO, Green A. The importance of genes and environment for ocular refraction and its determiners: a population based study among 20-45 year old twins. Br J Ophthalmol. 2001;85(12):1470–6.

47. Hammond CJ, Snieder H, Gilbert CE, Spector TD. Genes and environment in refractive error: the twin eye study. Invest Ophthalmol Vis Sci. 2001;42(6):1232–6.

48. Teikari JM, Kaprio J, Koskenvuo MK, Vannas A. Heritability estimate for refractive errors--a population-based sample of adult twins. Genet Epidemiol. 1988;5(3):171–81.

49. Dirani M, Chamberlain M, Shekar SN, Islam AF, Garoufalis P, Chen CY, et al. Heritability of refractive error and ocular biometrics: the Genes in Myopia (GEM) twin study. Invest Ophthalmol Vis Sci. 2006;47(11):4756–61.

50. Lopes MC, Andrew T, Carbonaro F, Spector TD, Hammond CJ. Estimating heritability and shared environmental effects for refractive error in twin and family studies. Invest Ophthalmol Vis Sci. 2009;50(1):126–31.

51. Verhoeven VJ, Hysi PG, Wojciechowski R, Fan Q, Guggenheim JA, Hohn R, et al. Genome-wide meta-analyses of multiancestry cohorts identify multiple new susceptibility loci for refractive error and myopia. Nat Genet. 2013;45(3):314–8.

52. Kiefer AK, Tung JY, Do CB, Hinds DA, Mountain JL, Francke U, et al. Genome-wide analysis points to roles for extracellular matrix remodeling, the visual cycle, and neuronal development in myopia. PLoS Genet. 2013;9(2):e1003299.

53. Flitcroft DI, Loughman J, Wildsoet CF, Williams C, Guggenheim JA, for the CC. Novel Myopia Genes and Pathways Identified From Syndromic Forms of Myopia. Invest Ophthalmol Vis Sci. 2018;59(1):338–48.

54. Tedja MS, Wojciechowski R, Hysi PG, Eriksson N, Furlotte NA, Verhoeven VJM, et al. Genome-wide association meta-analysis highlights light-induced signaling as a driver for refractive error. Nat Genet. 2018;50(6):834–48.

55. McGlinn AM, Baldwin DA, Tobias JW, Budak MT, Khurana TS, Stone RA. Form-deprivation myopia in chick induces limited changes in retinal gene expression. Invest Ophthalmol Vis Sci. 2007;48(8):3430–6.

56. Brand C, Schaeffel F, Feldkaemper MP. A microarray analysis of retinal transcripts that are controlled by image contrast in mice. Mol Vis. 2007;13:920–32.

57. Shelton L, Troilo D, Lerner MR, Gusev Y, Brackett DJ, Rada JS. Microarray analysis of choroid/RPE gene expression in marmoset eyes undergoing changes in ocular growth and refraction. Mol Vis. 2008;14:1465–79.

58. Schippert R, Schaeffel F, Feldkaemper MP. Microarray analysis of retinal gene expression in chicks during imposed myopic defocus. Mol Vis. 2008;14:1589–99.

59. Frost MR, Norton TT. Alterations in protein expression in tree shrew sclera during development of lens-induced myopia and recovery. Invest Ophthalmol Vis Sci. 2012;53(1):322–36.

60. Gao H, Frost MR, Siegwart JT, Jr., Norton TT. Patterns of mRNA and protein expression during minus-lens compensation and recovery in tree shrew sclera. Mol Vis. 2011;17:903–19.

61. Siegwart JT, Jr., Norton TT. The time course of changes in mRNA levels in tree shrew sclera during induced myopia and recovery. Invest Ophthalmol Vis Sci. 2002;43(7):2067–75.

62. Riddell N, Giummarra L, Hall NE, Crewther SG. Bidirectional Expression of Metabolic, Structural, and Immune Pathways in Early Myopia and Hyperopia. Frontiers in neuroscience. 2016;10:390.

63. Riddell N, Crewther SG. Integrated Comparison of GWAS, Transcriptome, and Proteomics Studies Highlights Similarities in the Biological Basis of Animal and Human Myopia. Invest Ophthalmol Vis Sci. 2017;58(1):660–9.

64. Tkatchenko TV, Shen Y, Tkatchenko AV. Analysis of postnatal eye development in the mouse with high-resolution small animal magnetic resonance imaging. Invest Ophthalmol Vis Sci. 2010;51(1):21–7.

65. Tkatchenko TV, Tkatchenko AV. Ketamine-xylazine anesthesia causes hyperopic refractive shift in mice. J Neurosci Methods. 2010;193(1):67–71.

66. Tkatchenko TV, Shen Y, Tkatchenko AV. Mouse experimental myopia has features of primate myopia. Invest Ophthalmol Vis Sci. 2010;51(3):1297–303.

67. Tkatchenko TV, Shen Y, Braun RD, Bawa G, Kumar P, Avrutsky I, et al. Photopic visual input is necessary for emmetropization in mice. Exp Eye Res. 2013;115C:87–95.

68. Storey JD, Tibshirani R. Statistical significance for genomewide studies. Proc Natl Acad Sci U S A. 2003;100(16):9440–5.

69. Huang da W, Sherman BT, Lempicki RA. Systematic and integrative analysis of large gene lists using DAVID bioinformatics resources. Nat Protoc. 2009;4(1):44–57.

70. Walter W, Sanchez-Cabo F, Ricote M. GOplot: an R package for visually combining expression data with functional analysis. Bioinformatics. 2015;31(17):2912–4.

71. Shah RL, Guggenheim JA, Eye UKB, Vision C. Genome-wide association studies for corneal and refractive astigmatism in UK Biobank demonstrate a shared role for myopia susceptibility loci. Hum Genet. 2018;137(11-12):881–96.

72. de Leeuw CA, Mooij JM, Heskes T, Posthuma D. MAGMA: generalized gene-set analysis of GWAS data. PLoS Comput Biol. 2015;11(4):e1004219.

73. Churchill GA, Airey DC, Allayee H, Angel JM, Attie AD, Beatty J, et al. The Collaborative Cross, a community resource for the genetic analysis of complex traits. Nat Genet. 2004;36(11):1133–7.

74. Tedja MS, Haarman AEG, Meester-Smoor MA, Kaprio J, Mackey DA, Guggenheim JA, et al. IMI - Myopia Genetics Report. Invest Ophthalmol Vis Sci. 2019;60(3):M89–M105.

75. Tkatchenko TV, Troilo D, Benavente-Perez A, Tkatchenko AV. Gene expression in response to optical defocus of opposite signs reveals bidirectional mechanism of visually guided eye growth. PLoS Biol. 2018;16(10):e2006021.

76. Riddell N, Crewther SG. Novel evidence for complement system activation in chick myopia and hyperopia models: a meta-analysis of transcriptome datasets. Scientific reports. 2017;7(1):9719.

77. Stone RA, McGlinn AM, Baldwin DA, Tobias JW, Iuvone PM, Khurana TS. Image defocus and altered retinal gene expression in chick: clues to the pathogenesis of ametropia. Invest Ophthalmol Vis Sci. 2011;52(8):5765–77.

78. Ronquillo CC, Bernstein PS, Baehr W. Senior-Loken syndrome: a syndromic form of retinal dystrophy associated with nephronophthisis. Vision Res. 2012;75:88–97.

79. Marmorstein LY, Munier FL, Arsenijevic Y, Schorderet DF, McLaughlin PJ, Chung D, et al. Aberrant accumulation of EFEMP1 underlies drusen formation in Malattia Leventinese and age-related macular degeneration. Proc Natl Acad Sci U S A. 2002;99(20):13067–72.

80. Tuschl K, Clayton PT, Gospe SM, Jr., Gulab S, Ibrahim S, Singhi P, et al. Syndrome of hepatic cirrhosis, dystonia, polycythemia, and hypermanganesemia caused by mutations in SLC30A10, a manganese transporter in man. Am J Hum Genet. 2012;90(3):457–66.

81. Leyva-Illades D, Chen P, Zogzas CE, Hutchens S, Mercado JM, Swaim CD, et al. SLC30A10 is a cell surface-localized manganese efflux transporter, and parkinsonism-causing mutations block its intracellular trafficking and efflux activity. J Neurosci. 2014;34(42):14079–95.

82. Papal S, Monti CE, Tennison ME, Swaroop A. Molecular dissection of cone photoreceptor-enriched genes encoding transmembrane and secretory proteins. J Neurosci Res. 2019;97(1):16–28.

83. Paridaen JT, Janson E, Utami KH, Pereboom TC, Essers PB, van Rooijen C, et al. The nucleolar GTP-binding proteins Gnl2 and nucleostemin are required for retinal neurogenesis in developing zebrafish. Dev Biol. 2011;355(2):286–301.

84. MacDonald R, Barbat-Artigas S, Cho C, Peng H, Shang J, Moustaine A, et al. A Novel Egr-1-Agrin Pathway and Potential Implications for Regulation of Synaptic Physiology and Homeostasis at the Neuromuscular Junction. Front Aging Neurosci. 2017;9:258.

85. Mann S, Kroger S. Agrin is synthesized by retinal cells and colocalizes with gephyrin [corrected]. Mol Cell Neurosci. 1996;8(1):1–13.

